# Statins inhibit onco-dimerization of the 4Ig isoform of B7-H3

**DOI:** 10.1101/2024.12.18.628944

**Authors:** Margie N. Sutton, Sarah E. Glazer, Ajlan Al Zaki, Arianna Napoli, Ping Yang, Priya Bhosale, Jinsong Liu, Seth T. Gammon, David Piwnica-Worms

## Abstract

B7-H3 (CD276), a member of the B7-family of immune checkpoint proteins, has been shown to have immunological and non-immunological effects promoting tumorigenesis [1, 2] and expression correlates with poor prognosis for many solid tumors, including cervical, ovarian and breast cancers [3–6]. We recently identified a tumor-cell autochthonous tumorigenic role for dimerization of the 4Ig isoform of B7-H3 (4Ig-B7-H3) [7], where 4Ig-B7-H3 dimerization *in cis* activated tumor-intrinsic cellular proliferation and tumorigenesis pathways, providing a novel opportunity for therapeutic intervention. Herein, a live cell split-luciferase complementation strategy was used to visualize 4Ig-B7-H3 homodimerization in a high-throughput small molecule screen (HTS) to identify modulators of this protein-protein interaction (PPI). Notably, the HTS identified several compounds that converged on lipid metabolism (including HMG-CoA reductase inhibitors, also known as statins) as significant inhibitors of 4Ig-B7-H3 dimerization (p < 0.01). *In vitro* and *in vivo* murine studies provided evidence that statin-mediated disruption of 4Ig-B7-H3 dimerization was associated with anti-tumor effects. Statin-mediated anti-cancer efficacy was selective for B7-H3-expressing tumors and retrospective analysis of clinical tumor specimens supported the hypothesis that concurrent statin use enhanced clinical outcomes for patients in a B7-H3 restricted manner. Thus, disruption of 4Ig-B7-H3 dimerization provides an unanticipated molecular mechanism linking statin use in cancer therapy and prevention with immune checkpoint.

## Introduction

The B7 family of immunoglobulins consists of structurally related, cell-surface protein ligands that can engage CD28 family receptors on lymphocytes to produce ‘costimulatory’ or ‘coinhibitory’ signals to augment or attenuate, respectively, immune response [8, 9]. Understanding these protein-protein interactions (PPIs) has led to a new era in cancer treatment, where checkpoint therapy by monoclonal antibodies inhibiting CTLA-4 and PD-1 on T-cells, or PD-L1 on tumor cells has revolutionized cancer therapy, shaping the landscape of cancer immunotherapy [10, 11] [12]. Many of the B7 family members, including PD-L1, PD-L2, B7-H3, B7-H4 and HHLA-2, can also be expressed on many nonhematopoietic cells and tumors, but the temporal and spatial differences in ligand expression and their functional contributions to pathogenic and protective immune responses have not been fully elucidated. In particular, B7-H3 (CD276), has garnered significant attention in recent years as a potential immune checkpoint molecule, given that the protein is widely overexpressed on tumor cell surfaces and vascular endothelium [5, 6, 13] and restricted expression on normal tissues. Many studies have begun to probe the tumor-intrinsic oncogenic role of B7-H3 tumor expression, as well as its role in tumor immunity. Studies have directly linked tumor-expressed B7-H3 protein with increased drug resistance [14], promotion of metastasis [2], and increased proliferation, invasion and migration [1]. In addition, correlative studies have documented a clear relationship between overexpression of B7-H3 on the tumor and poor prognosis for many solid tumors, including gynecological cancers such as cervical, ovarian and breast cancers [3–6]. However, the cognate receptor(s) has not been identified and the role of B7-H3 in immune response and tumor evasion has not been fully elucidated [15]. Interestingly, B7-H3 is the only B7 family member that contains a gene duplication resulting in an extracellular domain (ECD) containing a paired set of Igs (4Ig), whereas most other family members contain a common ECD comprising of a variable and constant Ig pair (2Ig isoform). Recently, quantitative molecular imaging techniques were used to interrogate the nanoscale localization of 4Ig-B7-H3 isoforms on tumor cells, identifying a previously unappreciated tumorigenic role for 4Ig-B7-H3 homodimerization [7].

Ongoing targeted drug development has leveraged tumor cell-expression of B7-H3 for therapeutic intervention, including antibody-based drugs [15] and drug-conjugates [16], CAR-based therapies [17, 18], and radio-ligand therapy [19, 20]. However, considering our recent findings that revealed a dimerization-dependent tumorigenic *cis* signaling role for 4Ig-B7-H3, we set out to identify small molecule inhibitors of this protein-protein interaction that could be quickly translated for clinical utility. Utilizing a live-cell luciferase complementation assay amenable to high throughput screening (HTS), we identified several compounds that converged on lipid metabolism, including HMG-CoA reductase inhibitors, and in particular statins, as significant inhibitors of 4Ig-B7-H3 dimerization (p < 0.01). *In vitro* and *in vivo* studies provided evidence that disruption of 4Ig-B7-H3 dimerization was associated with anti-tumor activity. Herein, statin-mediated anti-cancer efficacy was dependent upon on B7-H3 tumor expression and retrospective analysis of clinical tumor specimens supported the hypothesis that the well documented enhanced clinical outcomes for patients using statins are mechanistically linked to B7-H3 expression in the tumors. This study thus identifies a cholesterol-triggered pathway regulating tumorigenesis and indicates that interventions with cholesterol-lowering agents may complement current prevention and treatment strategies for patients with B7-H3-expressing tumors.

## Results

### A high-throughput screen identifies modulators of B7-H3 dimerization/multimerization

In the context of studies to dissect the mechanism(s) of 4Ig-B7-H3 biochemical functions and activation, a doxycycline-inducible split-luciferase complementation assay to monitor 4Ig-B7-H3 proximity and multimerization in real time [7] was developed using the ReBiL2.0 platform [21, 22]. This luminescence platform was used to conduct a plate-reader based high-throughput small molecule screen to identify modulators of 4Ig-B7-H3 dimerization/multimerization (from now on referred to as dimerization as it is the smallest component protein-protein interaction) in both directions (increasing or inhibiting) following the addition of a compound library **(Figure 1A)**. Assay screening conditions were determined in a step-wise fashion using our positive and negative controls and Z’ as criteria to evaluate each component of the assay setup and design. Optimal screening conditions were determined, which included a 384-well plate format, pre-plating of the cells with or without doxycycline (to induce 4Ig-B7-H3 expression), and addition of D-luciferin for assay readout **(Methods and Supplemental Figure 1, Supplemental Table 4)**. In addition, inter- and intra-assay variation was assessed using high, medium, and low concentrations of doxycycline to titrate 4Ig-B7-H3 protein expression levels, and thus dimerization potential of 4Ig-B7-H3. Using an interleaved-signal format, coefficients of variation were calculated for each signal (Max, Min and Mid) on each plate and all were found to be less than 20%. Signal windows and Z’ factors for each plate were ≥ 2 or ≥ 0.4, respectively, meeting standard criteria of acceptance. Visually, the assay was devoid of edge effects or drift **(Supplemental Figure 1C),** consistent with a robust platform readily available for a high throughput screen.

**Figure 1.**
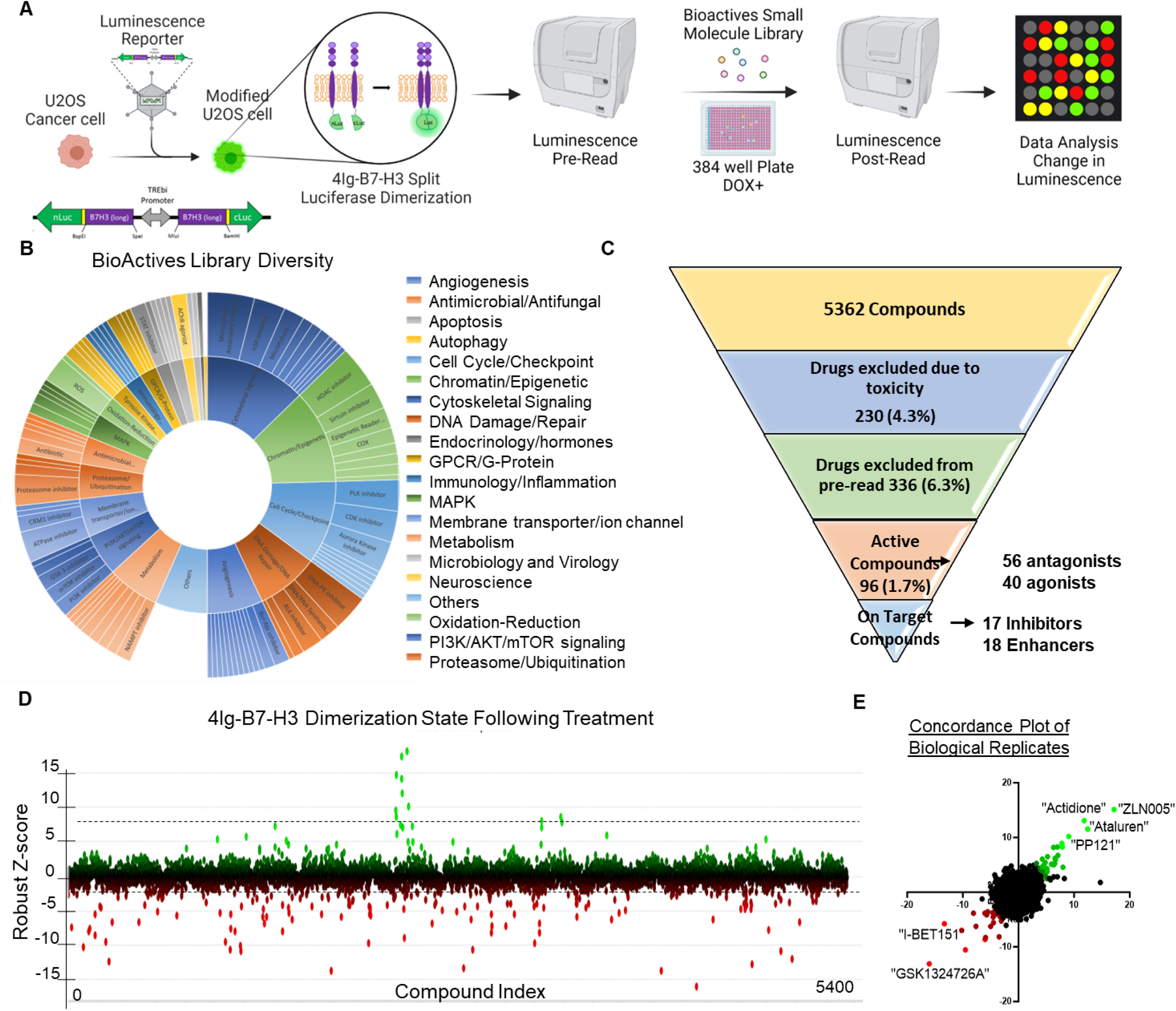
High-throughput screen identifies modulators of B7-H3 dimerization/multimerization. **A.** Schematic of the screening process is outlined. Briefly, U2OS cells were generated to stably express the 4IgB7-H3 split-luciferase construct. The cells were seeded and luminescence was pre-read prior to the addition of the small molecules. Luminescence was re-read and the change in relative luminescence was calculated to determine modulators of B7-H3 dimerization/multimerization. **B.** The small molecule library used for the high throughput screen comprised of 5,362 compounds targeting a wide range of proteins involved in cell function, or with known clinical benefit. Sunburst visualization of validated protein targets of the small molecules in the set using protein family hierarchy. 936 compounds within the library do not have validated protein targets (other) but they were included because they affect a known process or pathway. **C.** Hierarchical narrowing of hits obtained through the screening process following removal of toxic compounds and following counter screens to eliminate chemical and biological cofounders. 1.7% of the full library were active in our initial bioluminescence assay of dimerization/multimerization and of these 96 compounds, 35 (36.4%) were specific to modulating B7-H3 dimerization/multimerization compared with reporter inhibitors or changes to membrane associated proteins broadly. **D.** Robust Z-score on a per-compound basis was used to determine the degree of B7-H3 dimerization/multimerization modulation, where green or positive Robust-Z scores represented an increase in dimerization, and red or negative Robust-Z scores represented a decrease or inhibition of dimerization. Dotted lines represent the pre-defined threshold (99% CI) for activity. **E.** Using the robust Z-score per-compound between two biological replicates of the screen, the concordance was plotted where enhancers of 4Ig-B7-H3 dimerization were noted in green and inhibitors in red.

### Library Composition and Assay Development

The high throughput screening library (Bioactives Compound Library, Texas A&M IBT Drug Screening Core) used consisted of 5362 compounds with defined biological activity. The library included compounds that affect most cellular processes and drug target classes including: GPCR ligands, second messenger modulators, nuclear receptor ligands, actin and tubulin modulators, kinase inhibitors, protease inhibitors, ion channel blockers, gene regulation agents, lipid biosynthesis inhibitors, phosphodiesterase inhibitors as well as many others providing a diverse tool set for mechanism profiling, or lead compound screening **(Figure 1B)**. The screen was performed in biological duplicate, where the Robust Z and Z’ scores maintained consistency among all 18 plates in both replicates **(Supplemental Figure 2),** followed by the counter screen to remove non-specific toxic compounds as well as those that modulated split-luciferase interaction not specific to B7-H3 multimerization/dimerization, resulting in 35 ‘on target’ compounds consisting of 17 selective inhibitors and 18 selective enhancers **(Figure 1C)**. The Robust Z-score was determined for each compound based on the change in luminescence between the pre-read and post-read of the assay at 4 hours incubation and was plotted by each compound **(Figure 1D)**, where compounds in green enhanced B7-H3 dimerization and those in red reduced B7-H3 dimerization. The Robust Z-score was then plotted as a concordance plot (R squared = 0.1238, p< 0.0001) between both biological replicates of the screen, again documenting the reproducible enhancers in green and inhibitors in red **(Figure 1E)**.

### High-Throughput Screen HITs: Convergence on lipid metabolism as a key disruptor of 4Ig-B7-H3 dimerization

To eliminate biological and chemical confounders, secondary screens were employed. CellTiter-Glo was used to measure ATP levels as an endpoint toxicity assay in parental cells with and without doxycycline addition (to induce B7-H3 expression), thus, providing an opportunity to eliminate generally toxic compounds and identify potential B7-H3-dependent toxic inhibitors that were not previously identified as modulators of B7-H3 dimerization in the initial 4-hour dimerization window. In addition, active compounds were further narrowed by performing a counter screen using our previously published K-Ras: K-Ras dimerization cell line [22, 23]. Due to the isogenic nature of this system, using this cell line as a counter screen allowed for elimination of compounds that were not selective to B7-H3 dimerization (e.g., those that interfered with cellular ATP levels (necessary for the luciferase enzymatic reaction), those that disrupted split-luciferase complementation, those that directly modulated firefly luciferase, or those that altered U2OS transcription/translation/degradation) **(Figure 2A)**. Target pathway analysis was performed for each group of selective inhibitors and enhancers and visualized using a sunburst plot **(Figure 2B-C)**. Inhibitors targeting the PI3K/AKT/mTOR signaling pathway, cell cycle, PKC and JAK kinases, epigenetic regulation, such as bromodomain inhibitors, and lipid metabolism/cholesterol lowering compounds were inhibitors of 4Ig-B7-H3 dimerization. Interestingly, several other PI3K/AKT/mTOR signaling inhibitors (specifically mTORi) were capable of enhancing 4Ig-B7-H3 dimerization, as were PGC1α inhibitors, and eIF4G1 inhibitors, but most 4Ig-B7-H3 enhancers did not belong to a single cellular process or class of molecules. The ‘active hits’ were then further subjected to a 3-dose titration curve **(Supplemental Figure 3)** and further validated using bioluminescence live cell imaging with an IVIS100. Treatment of HMG-CoA reductase inhibitors (statins; Mevastatin and Atorvastatin Caclium (1-100 nM) significantly inhibited B7-H3 multimerization **(Figure 2D-E)**. To assess the functional consequence of inhibiting 4Ig-B7-H3 dimerization, clonogenic assays were performed in the presence (DOX+) and absence (DOX-) of 4Ig-B7-H3. Statins significantly inhibited clonogenic growth in the presence of B7-H3 expression (DOX+) **(Figure 2F-G, Supplemental Figure 4),** where the relative inhibition compared to diluent treated control without doxycycline expression was less than 5% whereas the relative change in long-term clonogenic growth with statin treatment in B7-H3 expressing (DOX+) cells was 25% **(Figure 2F)**. This B7-H3 dependence in growth inhibition was maintained using HeLa WT and paired Crispr/Cas9 KO cells [7], and SKOv3-ip and MDA-MB-231 cells which endogenously express B7-H3, where the relative reduction in clonogenic grown following statin treatment in B7-H3 expressing cells was 25-70% **(Supplemental Figure 4)**. HMG-CoA reductase inhibitors or statins inhibit the rate limiting step in the mevalonate pathway leading to the production of cholesterol and are routinely used clinically to lower blood cholesterol levels and reduce risk for illnesses related to atherosclerosis. Interestingly, while several inhibitors of B7-H3 dimerization did not converge on a signal signaling pathway, there was convergence on pathways that alter lipid/cholesterol metabolism.

**Figure 2.**
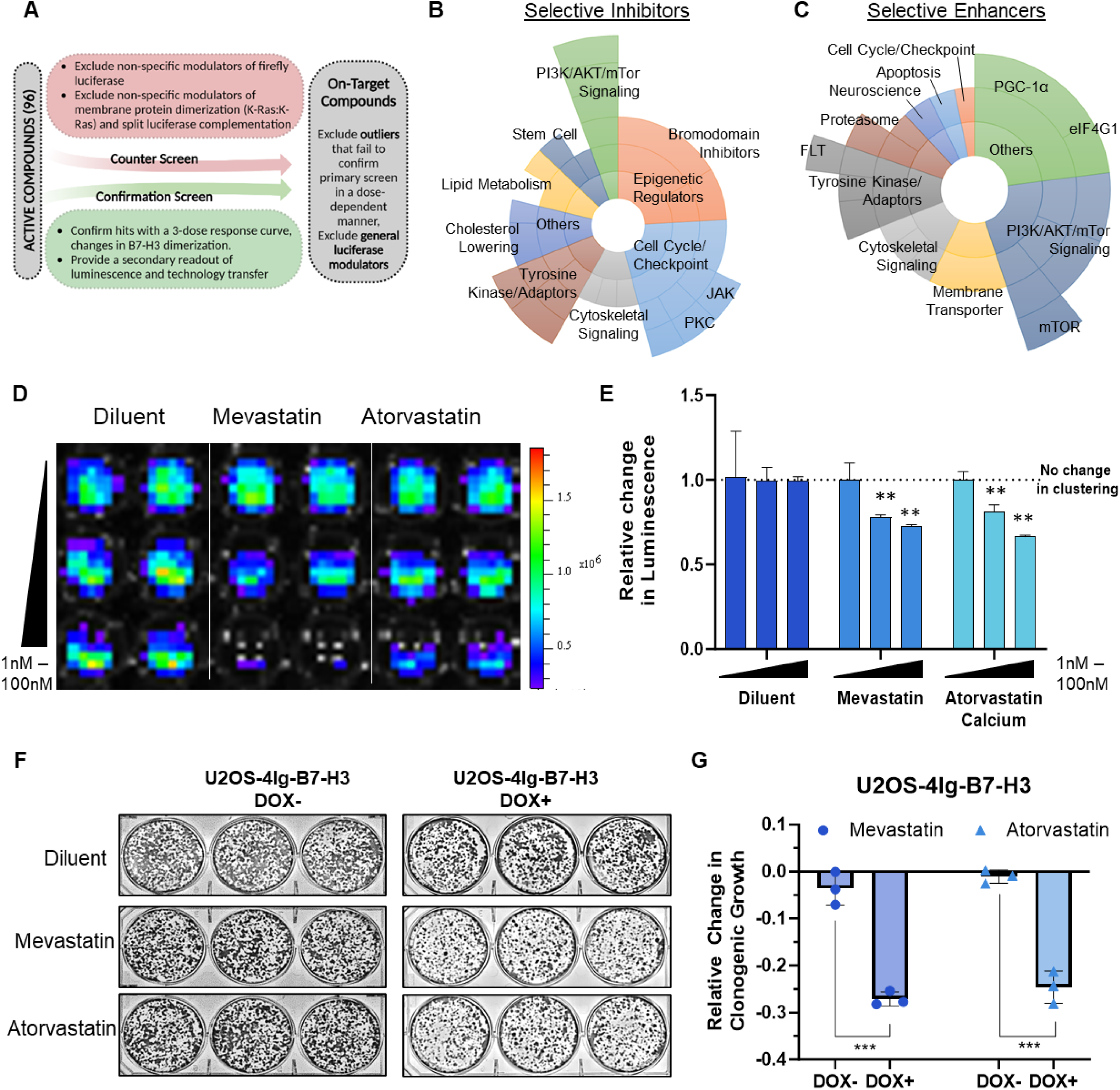
Statins significantly inhibit 4Ig-B7-H3 dimerization. **A.** Schematic of the screening process used to define “On-target” compounds as opposed to active compounds is outlined. Both counter screens and confirmation screens were used to exclude outliers that failed to confirm primary screening data in a dose-dependent manner as well as general luciferase modulators. **B.** Sunburst visualization of validated selective “on-target (4Ig-B7-H3-dimerization)” inhibitors based on their pathway or protein targets of the small molecules in the set using protein family hierarchy. **C.** Sunburst visualization of validated selective “on-target (4Ig-B7-H3-dimerization)” enhancers based on their pathway or protein targets of the small molecules in the set using protein family hierarchy. **D.** U2OS-4IgB7-H3 cells were seeded in 96-well black-walled plates and incubated with or without doxycycline for 48 hours and 200 nM D-Luciferin. Cells were treated with compounds as indicated and bioluminescent signals were measured 4 hours post addition using an IVIS100 imaging system. Bioluminescence signal in doxycycline-induced U2OS-4Ig-B7-H3 ReBiL cells was significantly higher than in uninduced cells or empty split-Luc cells demonstrating 4Ig-B7-H3 dimerization/multimerization, and inhibition of dimerization was observed in a dose dependent manner for the tested compounds. **E.** Photon flux was calculated over three independent experiments performed in technical duplicate and represented in the bar graph (mean ± SD). Photon flux was significantly reduced (p<0.001) upon treatment with mevastatin, atorvastatin (0-100 nM). **F.** 5,000 U2OS-4Ig-B7-H3 cells were seeded and treated with or without doxycycline (500 µg/mL) every 3 days with and without statin treatment (1µM treatment with mevastatin, atorvastatin calcium), on day 1 and day 4 and clonogenic growth was measured after 10 days. Colonies were fixed and stained with Coomassie blue on day 10. **G.** Clonogenic growth was quantified using ImageJ and represented in the bar graph (mean ± SD). The experiment was performed in technical triplicate with three biological replicates, ***p<0.001.

### B7-H3 co-localizes with cholesterol in the plasma membrane and B7-H3 multimerization is affected by changing cellular cholesterol levels or cholesterol binding

These findings led us to investigate the link between lipid/cholesterol metabolism (statins regulated pathway) and B7-H3. It is well established that cholesterol can control the activity of a wide range of membrane receptors through several mechanisms including membrane fluidity, curvature, and more directly by binding to the membrane-spanning domains of susceptible receptors to modulate their conformation and dimerization state (e.g., G-protein coupled receptors) [24]. Preliminary studies to identify possible mechanisms of convergence between decreased multimerization of 4Ig-B7-H3 and altering lipid/cholesterol metabolism revealed interesting structural features of the 4Ig-B7-H3 transmembrane domain that might lend itself to cholesterol binding. Using computational and modeling methods (SWISS-MODEL, [25, 26]), the protein sequence of 4Ig-B7-H3 was assessed for cholesterol binding consensus domains (CRAC/CARC) located near the outer and inner leaflet of the transmembrane domain (TMD). A cholesterol binding CRAC domain near the inner-leaflet of the TMD of 4Ig-B7-H3 was identified **(Figure 3A)**. Interestingly, analysis shows that many members of the B7-family of immune modulatory proteins also contain CRAC/CARC domains **(Supplementary Figure 5)**. Next, immunofluorescence staining was performed to determine the co-localization between cholesterol using Filipin III and endogenous B7-H3 using a high affinity monoclonal antibody **(Figure 3B)**. Pixel-wise correlation analysis showed significant co-localization between B7-H3 (green) and cholesterol (blue) as determined by Pearson’s correlation coefficient (R^2^=0.89), Mander’s split coefficient (tM1=0.795, tM2=0.815), and Spearman’s rank correlation coefficient (0.87). To interrogate the impact of cholesterol binding on 4Ig-B7-H3 dimerization states, hetero-FRET-FLIM (Forester Resonance Energy Transfer-Fluorescence Lifetime Imaging Microscopy) was utilized as previously described [7]. Hetero-FRET-FLIM was performed by measuring fluorescence lifetimes of expressed human 4Ig-B7-H3 fluorescently-tagged WT 4Ig-B7-H3-meGFP and WT 4Ig-B7-H3-mCherry on the HeLa *CD276^-/-^* KO background, wherein 4Ig-B7-H3 protein fusions were only expressed exogenously. By means of ultra-sensitive time-correlated single-photon counting (TCSPC) in the meGFP spectral window, we measured the fluorescence decays of meGFP-tagged 4Ig-B7-H3 in fixed cells to determine protein proximity and the potential for homo- and hetero-dimeric complexes **(Figure 3C)**. Upon decay correction and lifetime fitting, we found that a two compartment model best fit the lifetime decay curve and centered around our previously observed monomeric (2.6 ns) and dimeric states (1.3 ns) in a 1:2 monomeric/dimeric ratio **(Figure 3D)** [7]. Co-transfection HeLa *CD276^-/-^* cells with WT 4Ig-B7-H3-meGFP and Vector-mCherry resulted in a split lifetime abundance of 43/56 monomeric/dimeric ratio **(Figure 3D)**, both values similar to the abundance ratios observed previously with homo-FRET-FLIM [7]. Next a CRAC mutant (F486A) construct was generated, where confocal fluorescence imaging using wildtype 4Ig-B7-H3-meGFP (green) and 4Ig-B7-H3^mCRAC^-mCherry (red) confirmed proper localization of the mutant construct **(Figure 3E)**. Using the co-transfection method described previously, we performed hetero-FRET-FLIM and measured the meGFP lifetimes and their associated populations **(Figure 3F)**. Upon co-transfection of the WT 4Ig-B7-H3 construct with the CRAC mutant 4Ig-B7-H3 construct we observed a shift from a more abundant dimeric state (1.3 ns) to a more abundant monomeric state (2.6 ns) where the population ratios nearly reversed compared to the WT/WT co-transfection **(Figure 3F-G)**. Using the pharmacological disruptor of lipid rafts and cholesterol extractor, methyl-β-cyclodextrin (MBCD) [27], measured lifetime abundance of monomeric or dimeric/multimeric proteins also shifted to a more monomeric state ratio upon MBCD treatment for 10 minutes compared to control diluent **(Figure 3H)**. These results were confirmed in a secondary assay of B7-H3 dimerization utilizing the ReBiL2.0 split-luciferase complementation assay in U2OS cells where 4Ig-B7-H3 multimerization was significantly decreased in a concentration- and time-dependent manner upon MBCD treatment **(Figure 3I)**. These data indicate that decreased membrane cholesterol levels or the inability of mutant CRAC 4Ig-B7-H3 to bind cholesterol inhibited homo-dimeric protein-protein interactions of 4Ig-B7-H3 in live cells.

**Figure 3.**
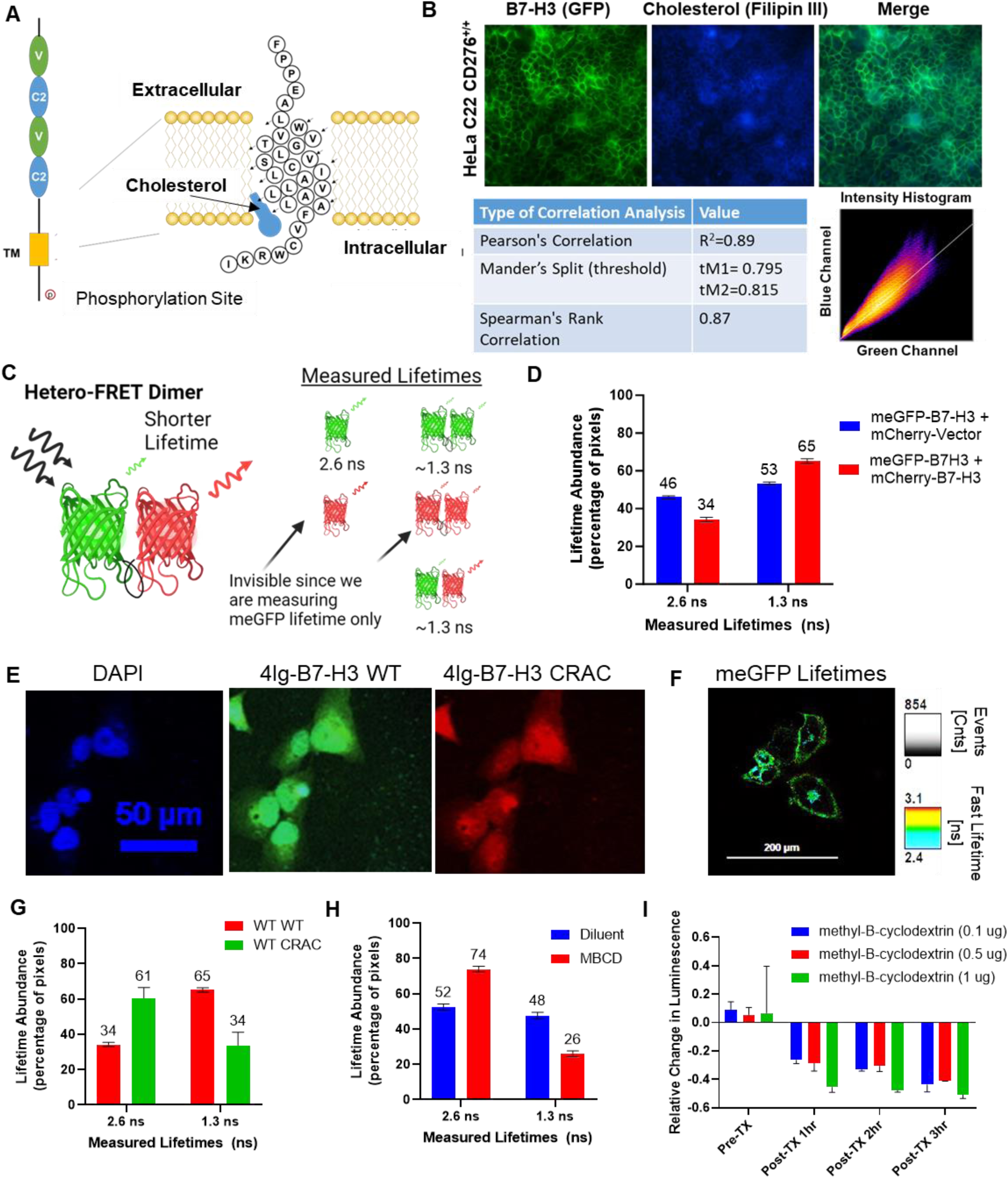
4Ig-B7-H3 dimerization is mediated through the CRAC/CARC domain and is cholesterol dependent. **A.** Schematic of the transmembrane domain and amino acid sequence of 4Ig-B7-H3 demonstrating the potential site of cholesterol binding at the predicted CRAC domain. **B.** Immunofluorescent staining of endogenous B7-H3 (green) and cholesterol (blue) in WT HeLa cervical cancer cells. Pixel wise correlation analysis showed significant correlation and co-localization between B7-H3 (green) and cholesterol (blue) as determined by Pearson’s Correlation (R^2^=0.89), Mander’s Split (tM1=0.795, tM2=0.815), and Spearman’s Rank Correlation (0.87). **C.** A scheme to demonstrate the potential monomer or dimeric fluorescent pairs that could be observed using the hetero-FRET-FLIM system. For example, monomeric GFP lifetime has a measured lifetime of 2.6 ns (monomer), compared to GFP lifetime measurements under homo-FRET conditions (dimeric state) that results in a reduced lifetime 1.3 ns, as energy transfer occurs between the two fluorescent proteins where their localization is less than 10 nm apart. Monomeric mCherry and homo-dimeric mCherry lifetimes are not observed in our experiment given that only GFP lifetimes were measured, however, hetero-dimeric complexes that contain GFP and mCherry would be observed under the experimental conditions, where the observed lifetime was approximately 1.3 ns (dimer). **D.** The average lifetime of GFP fluorescence determined for individual HeLa *CD276^-/-^* KO cells transiently transfected with *Vector-mCherry* and *CD276-mEGFP (V/CD276)* or *CD276-mmCherry* and *CD276-mEGFP (CD276/CD276)* and the relative abundance at monomeric (GFP alone) or dimeric (homo-or hetero-complexes) was measured using a bi-exponential fit. Bars represent the mean (± standard deviation) lifetimes measured for 10 individual cells over three biological replicates. **E.** Immunofluorescence microscopy was performed on HeLa CD276^-/-^ knockout cells re-expressing exogenous WT CD276-GFP and CRAC mutant CD276-mCherry. DAPI (nucleus), GFP (exogenous WT 4Ig-B7-H3) and mCherry (exogenous CRAC^Mut^4Ig-B7-H3) at 24 hours post transfection. Expected co-localization between the constructs was observed. Scale bars, 50 µm. **F.** Representative images showing GFP fluorescence lifetime (pseudocolored) of exogenously expressed C-terminally linked to WT *CD276* (4Ig-B7-H3-meGFP) co-transfected with WT mCherry-CD276. **G.** HeLa CD276^-/-^ cells co-transfected with WT CD276-meGFP and WT CD276-mCherry paired constructs or WT CD276-meGFP and CRAC^Mut^ CD276-mCherry. The relative abundance at monomeric (2.6 ns) or dimeric (homo- or hetero-complexes, 1.3 ns) was measured using a bi-exponential fit. CRAC^Mut^ CD276-mCherry reduced the abundance of dimeric complexes from 65% to 34%. Bars represent the mean (± standard deviation) lifetimes measured for 10 individual cells over three biological replicates. **H.** Similarly, HeLa CD276^-/-^ cells were transfected with WT CD276-meGFP and treated with or without MBCD (0.1 μg) for 10 minutes before washing and fixation. Lifetime abundance was measured and treatment with MBCD reduced the dimeric abundance from 48% to 26%. Bars represent the mean (± standard deviation) lifetimes measured for 10 individual cells over three biological replicates. **I.** U2OS-4IgB7-H3 cells were seeded in 96-well black-walled plates and incubated with or without doxycycline for 48 hours. Cells were treated (TX) with MBCD or diluent as indicated for 10 minutes followed by washing, the addition of 200 nM D-Luciferin, and then bioluminescent signals were measured hourly using an IVIS100 imaging system. Photon flux was calculated over three independent experiments performed in technical duplicate and represented in the bar graph as the relative change in luminescence compared to diluent-treated cells (mean ± SD).

### Inhibiting B7-H3 dimerization with statins reduces intrinsic oncogenic potential in a B7-H3 dependent fashion in vitro which may also influence the extrinsic tumor microenvironment

To determine the functional significance of inhibiting 4Ig-B7-H3 multimerization with statins, our model cell lines were used to test the effect of statin treatment on clonogenic growth. As previously reported [7], CRISPR/Cas9 was used to knockout *CD276* (B7-H3) expression in HeLa cervical cancer cells, where two stable cell pairs were generated and grown from a single clone to reduce the effect of clonal-specific cell line alterations, and then lentivirally transduced to re-express 4Ig-B7-H3 in a heterogeneous fashion [7]. In HeLa wildtype cells (C20, C22) that endogenously express 4Ig-B7-H3, treatment with mevastatin (1 µM) or atorvastatin (1 µM) significantly reduced clonogenic growth (>20%); however, upon knockout of *CD276* (C6, C11) treatment with statins did not significantly inhibit clonogenic growth compared to diluent control **(Figure 4A, Supplemental Figure 6A)**. Using the lentiviral re-expressed 4Ig-B7-H3 rescue HeLa cell lines, the statin-induced reduction of colony formation phenotype was restored **(Figure 4B, Supplemental Figure 6B)**. Thus, the effect of statin-mediated long term clonogenic colony formation was dependent upon 4Ig-B7-H3 expression. Similarly, the 4Ig-B7-H3 dependent phenotype of statin-inhibited clonogenic growth was observed in SKOv3-ip wildtype cells, which endogenously express 4Ig-B7-H3 (C20, **Figure 4C**; C10 **Supplemental Figure 6C**), but was not observed in the paired knockout cell lines (C11, C4). MDA-MB-231, triple negative breast cancer cells, which also endogenously express 4Ig-B7-H3 were sensitive to statin treatment, where significant changes in clonogenic growth (>60%) were observed **(Supplemental Figure 6D)**. ImageJ was used to quantify the relative change in clonogenic growth for all cell lines, for three independent experiments performed in technical triplicate **(Figure 4D-F, Supplemental Figure 6A-D)**. Endogenous expression of B7-H3 was confirmed using immunofluorescence staining of SKOv3-ip and MDA-MB-231 cells **(Supplemental Figure 6E)**.

**Figure 4.**
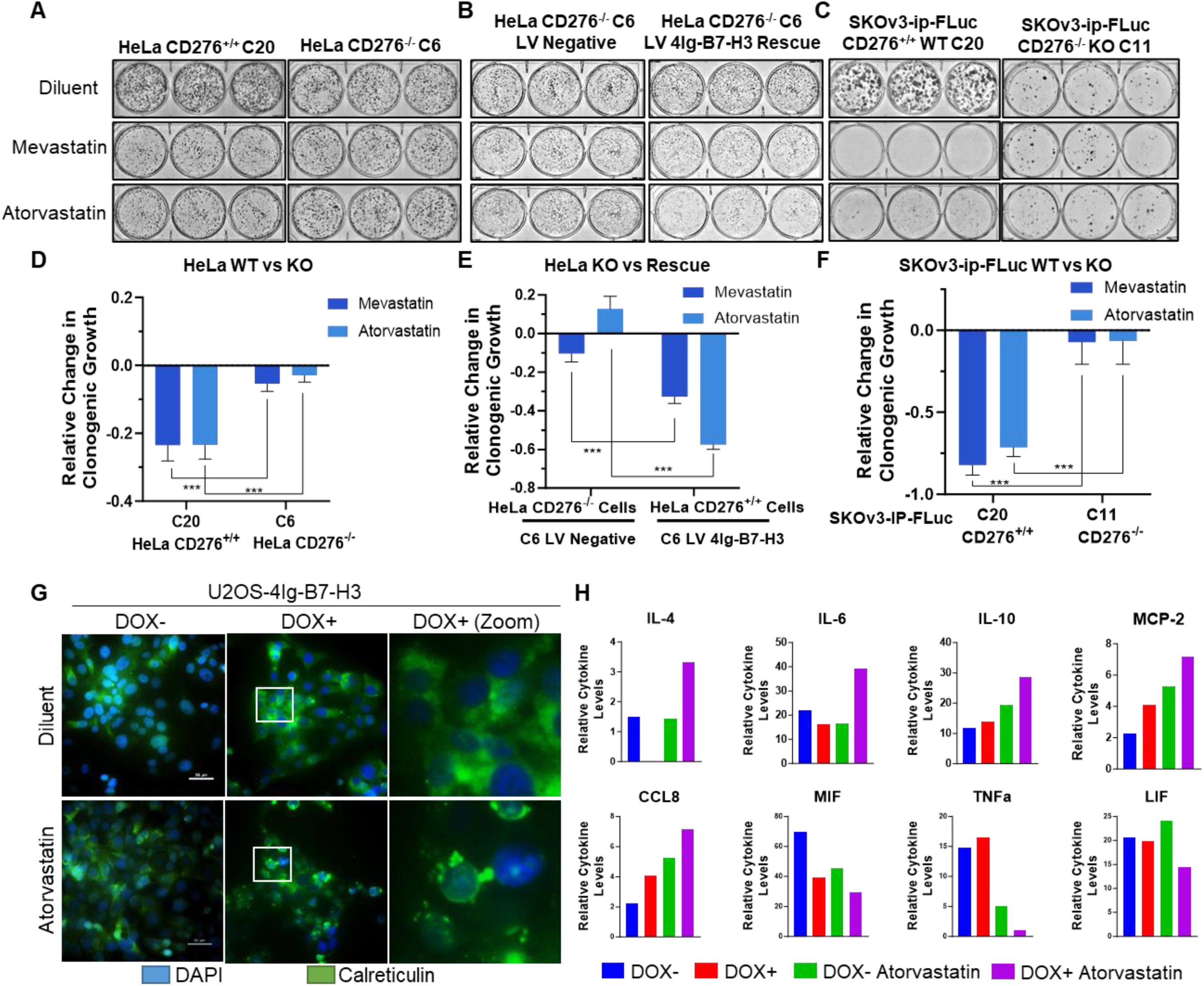
Inhibiting B7-H3 dimerization with statins reduces intrinsic oncogenic potential in a B7-H3 dependent fashion in vitro which may also influence the extrinsic tumor microenvironment. **A-B.** 2,000 HeLa wildtype or *CD276* knockout cells, or rescued cells were seeded and allowed to grow for 10 days, changing the media every 72 hours. Cells were subjected to 1µM treatment with mevastatin or atorvastatin calcium on day 1 and day 4. Plates were stained with Coomassie blue dye on day 10 and representative images of the plates are shown. **C.** 5,000 SKOv3-ip wildtype or *CD276* knockout cells were seeded and allowed to grow for 10 days, changing the media every 72 hours. Cells were subjected to 1µM treatment with mevastatin or atorvastatin calcium on day 1 and day 4. Plates were stained with Coomassie blue dye on day 10 and representative images of the plates are shown. **D-F.** Clonogenic growth was quantified using ImageJ and represented in the bar graph (mean ± SD). The experiment was performed in technical triplicate with three biological replicates, ***p<0.001. **G.** 1.0 x 105 U2OS-4Ig-B7-H3 cells were seeded in chamber slides and treated with or without Doxycycline for 48 hours. Cells were treated with diluent or atorvastatin (1 μM) for 24 hours prior to fixation and immunofluorescence staining of endogenous calreticulin. In the presence of doxycycline-induced 4Ig-B7-H3 expression and dimerization, atorvastatin treatment resulted in significant translocation and surface punctate expression of CALR. Scale bars; 50 μm **H.** Media obtained from the same experimental conditions as described in **(G)** was used to perform cytokine analysis. Relative cytokine levels were plotted for each condition; DOX-(blue), DOX+ (red), DOX-Atorvastatin treatment (green), DOX+ Atorvastatin treatment (purple). Significant changes in cytokine levels as determined by two-way ANOVA are plotted as relative change.

Given the tumor-expressed 4Ig-B7-H3 dependence of statins to significantly inhibit clonogenic growth in multiple models, we began to explore potential mechanisms of tumor cell death that could relate to the immune modulatory functions of 4Ig-B7-H3. Immunogenic cell death (ICD) is one form of cell death in which anticancer agents can induce clinically-relevant tumor-targeting immune responses [28]. While there are many associated extracellular signals or damage-associated molecular patterns (DAMPs) that contribute, their combined release favors the recruitment and activation of antigen-presenting cells [29]. One such ICD-associated DAMP includes the surface-exposed endoplasmic reticulum (ER) chaperone, calreticulin (CALR). Immunofluorescence staining revealed significant translocation and surface punctate expression of CALR following atorvastatin treatment (1 μM) in a 4Ig-B7-H3 dependent manner **(Figure 4G)**. In addition, measurement of extracellular cytokine release following statin treatment of U2OS-4Ig-B7-H3 cells in the presence or absence of doxycycline (+/- 4Ig-B7-H3 expression and dimerization) **(Supplemental Figure 7)** revealed significant changes in levels of IL-4, IL-6, IL-10, MCP-2, CCL8, MIF, TNFα, and LIF **(Figure 4H)**. These data suggest that statin-mediated inhibition of 4Ig-B7-H3 dimerization *in vitro* alters intrinsic signaling pathways in a 4Ig-B7-H3-dependent fashion, likely influencing oncogenic growth, and the extrinsic tumor microenvironment.

### In vivo statin-mediated inhibition of tumor outgrowth depends on 4Ig-B7-H3 expression in the tumor cell compartment

Using continual release osmotic pumps to reduce pharmacologic valleys and peaks of therapeutic dosing, we treated athymic nude mice bearing subcutaneous HeLa xenografts (wildtype 4Ig-B7-H3 and knockout tumors) with a literature established dose of 10 mg/kg/day atorvastatin calcium or diluent for 28 days [30–32]. Tumors were implanted at the same time as the animals underwent surgical implantation of the osmotic treatment delivery pumps. Tumor measurements, mouse weight, and overall survival were monitored for each group (4 groups, n=10 mice each) throughout the duration of the study. Signs of toxicity were not observed due to diluent or atorvastatin treatment **(Supplemental Figure 8)**, as body weight steadily increased weekly, only to slightly diverge when the tumor size of the diluent-treated mice was significantly larger than those in the treatment or knockout cohorts, and clinical assessment of the mice did not detect changes in normal behavior or appearances. Overall survival mimicked the previously observed change in survival upon knockout of tumor-expressed 4Ig-B7-H3 **(Figure 5A)**, where mice bearing HeLa xenografts that did not express 4Ig-B7-H3 had a significantly longer overall survival compared to mice bearing HeLa wildtype xenografts expressing 4Ig-B7-H3 [7]; p=0.035 Log-rank (Mantel-Cox) test **(Figure 5A)**. Treatment with atorvastatin further enhanced survival, in a 4Ig-B7-H3 dependent manner (dashed dark blue line). This added therapeutic benefit of statin inhibition was not observed when tumor-expressed 4Ig-B7-H3 was knocked out (dashed light blue line), p=0.6521. Individual growth curves for WT HeLa bearing mice in each treatment group are shown in **Figure 5B**. The tumor outgrowth inhibitory effect of statin treatment on two additional orthotopic models of human female cancers including MDA-MB-231 triple negative breast cancer, and SKOv3-IP high grade serous ovarian cancer, was also tested **(Figure 5C-E)**. In both cases, treatment with 10 mg/kg daily atorvastatin significantly increased survival (MDA-MB-231, **Figure 5D**) and reduced tumor burden (MDA-MB-231, **Figure 5C**; SKOV3-ip, **Figure 5E**). Tumor expression of B7-H3 was confirmed for all models by immunohistochemical staining upon endpoint **(Figure 5F)**. To understand the impact of 4Ig-B7-H3 tumor genotype on statin-mediated survival *in vivo*, hazard ratios were obtained using a matrix table containing genotype, treatment stratification and overall survival. Using a Cox regression model, where outcome of treatment with statins was evaluated based on the expression of 4Ig-B7-H3 in the tumor compartment, it was observed that statin treatment in the context of 4Ig-B7-H3 expression (genotype 1) had a significant effect (p=0.008) on survival of the mice, decreasing the risk to death by 1.3-fold **(Figure 5G)**. These data *in vivo* confirmed our previous findings *in vitro* that statin-mediated inhibition of tumor outgrowth depended upon tumor-expressed 4Ig-B7-H3, likely mediated by disruption of 4Ig-B7-H3 dimerization.

**Figure 5.**
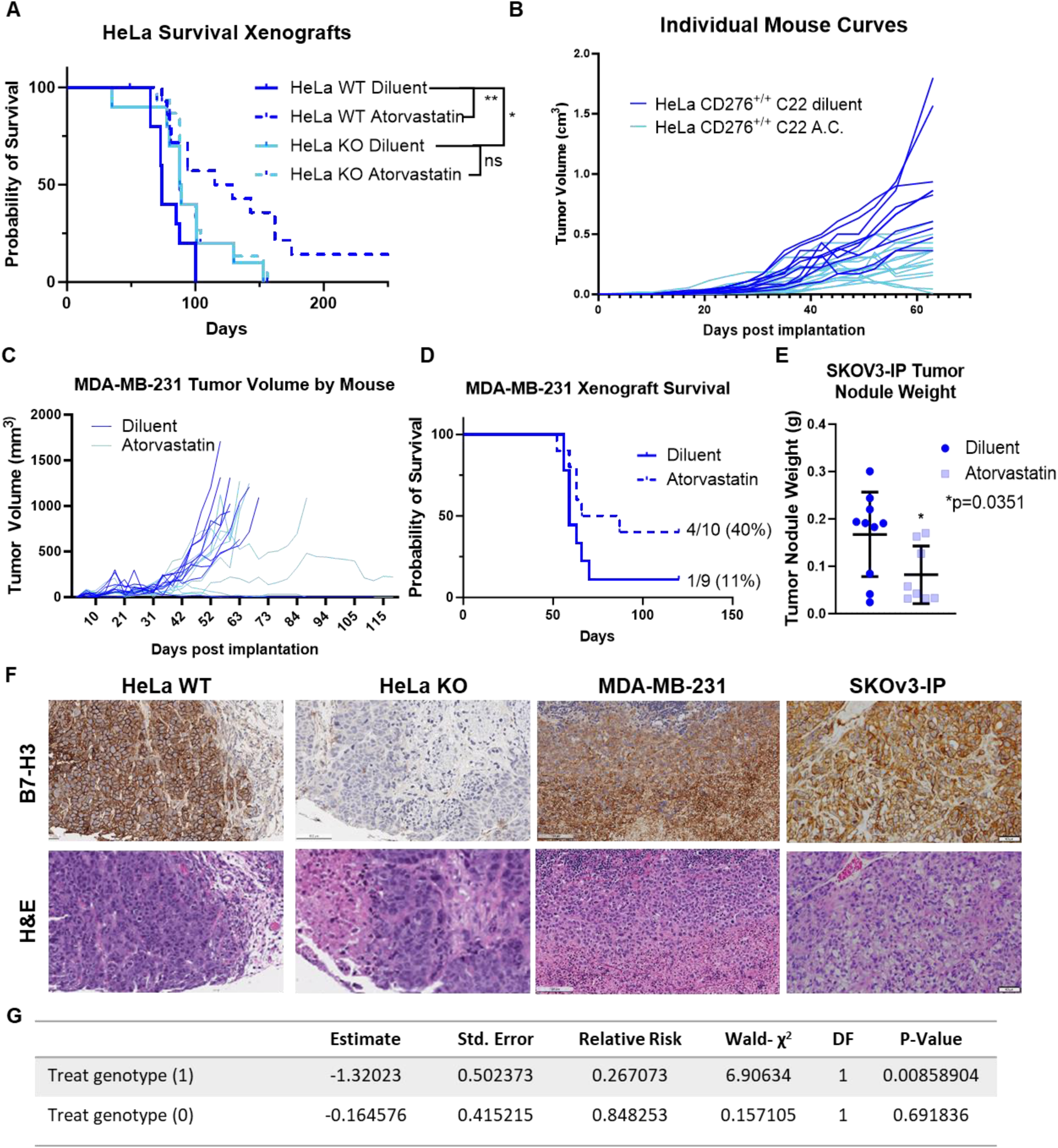
*In vivo* statin-mediated inhibition of tumor outgrowth depends on B7-H3 tumor expression in the tumor cell compartments. **A.** Kaplan-Meier plots of endpoint survival of HeLa WT and KO B7-H3 xenografts implanted on the right flank of 8 week old *nu/nu* mice treated with diluent or atorvastatin calcium (10 mg/kg). Statistically significant difference in survival between HeLa *CD276*^+/+^ (n=10) and HeLa *CD276*^-/-^ (n=10) was determined (p<0.035, Mantel-Cox Log-rank test). Treatment with atorvastatin further enhanced survival (HeLa WT dashed blue, p=0.005), in a B7-H3 dependent manner. This added therapeutic benefit of statin inhibition was not observed when tumor-expressed B7-H3 was knocked out (solid light blue vs. dashed light blue), p=0.6521. **B.** Tumor growth plotted independently by mouse for the diluent and atorvastatin treated WT B7-H3 expressing tumor xenografts. **C.** MDA-MB-231 xenografts were implanted orthotopically in the mammary fat pad of nu/nu 8 week old mice, and tumor volume was independently monitored by mouse for diluent or atorvastatin (10mg/kg) treated mice and endpoint survival was documented in a Kaplan-Meier plot **(D)**. Median survival for diluent treated mice was 59 days, where atorvastatin treated mice had an extended median survival of 76.5 days. **E.** SKOv3-ip tumor xenografts were grown orthotopically in the intraperitoneal cavity of 8 week old nu/nu mice. Tumor weight (g) following 4 weeks of atorvastatin treatment (10 mg/kg) or diluent was determined upon endpoint. Atorvastatin treatment significantly (p=0.035) reduced tumor weight. **F.** Endpoint histology was performed and representative H&E staining and human B7-H3 immunostaining is shown. Images captured with 4X objective, scale bars, 50 µm. **G.** Hazard ratios obtained from in vivo survival analysis data were used in a Cox regression model where outcome of treatment with statins was evaluated based on the expression of B7-H3 in the tumor compartment, it was observed that statin treatment in the context of B7-H3 expression (genotype 1) had a significant effect (p=0.008) on survival of the mice, decreasing the risk to death by 1.3-fold.

### Serum interferon-gamma, interlukin-6 and interleukin-10 serve as non-invasive pharmacodynamic biomarkers for prevention of tumor outgrowth with atorvastatin treatment

To identify biomarkers of response to statin treatment, in addition to tumor-expressed 4Ig-B7-H3, a receiver operator characteristic curve (ROC) was developed wherein prior studies were used to quantify the potential for a change in tumor volume to predict the fate of the mouse (responder/cure versus tumor progression) **(Figure 6A)**. By generating this ROC curve, mice were able to be humanely euthanized within each treatment arm (with or without tumor expressed 4Ig-B7-H3 (WT vs KO), with or without atorvastatin treatment (diluent vs. atorvastatin) after a sufficient change in tumor volume <0.03 mm^3^ over two measurements, collect sample specimens at this pre-designated the time of response, and compare to predicted endpoint outcomes to identify biomarkers of response to statin treatment and prevention of tumor outgrowth. Three to four mice per group were obtained, including potential cures (dark orange) and potential non-responders (light orange) in the WT tumor statin-treated group. Serum was analyzed for circulating cytokine changes, suspension CyTOF was performed on the spleens for immune cell profiling and relative abundance, and liver and tumor specimens were collected for immunohistochemistry and H&E staining **(Figure 6B, Supplemental Figure 9)**. Measuring a panel of 16 circulating cytokines involved in mouse innate immunity, we identified 3 non-invasive pharmacodynamic biomarkers of response to statin treatment. Non-responders had statistically significant higher levels of IFN-γ, IL-10, and IL-6 compared to those mice whose tumors were responding to statin treatment and likely to be cured **(Figure 6C)**. Significant changes in these cytokine levels were not detected in mice bearing WT or KO tumors treated with diluent, or mice bearing KO tumors treated with or without statins, reinforcing the interaction between statins and B7-H3-expression on the tumor **(Supplemental Figure 9)**. These circulating cytokines also affected immune cell populations within the spleen, where changes in a small subset of cell frequency was observed between the responders and non-responders depicted using a volcano plot of the -log(FDR) vs log(FC) **(Figure 6D)**. Namely, the frequency of the B-cell population was increased in the mice predicted to be cured by statin treatment **(Figure 6E)**, whereas an unassigned population (CM-unassigned) showed a significant decrease in population frequency in the responders **(Supplemental Figure 10)**. Interestingly, protein expression patterns between the B-cell groups were relatively similar between each individual responder and non-responder queried **(Figure 6F)**. Using a tissue microarray developed from formalin fixed paraffin embedded (FFPE) tissues from both the liver and tumor, immunohistochemical staining and H&E staining were performed. Significant changes in immune cell infiltration (F4/80 macrophages; CD11b myeloid; CD19 B-cells) did not reach a threshold of detection between responders versus non-responders for either the tumor or liver compartment **(Supplemental Figure 11)**.

**Figure 6.**
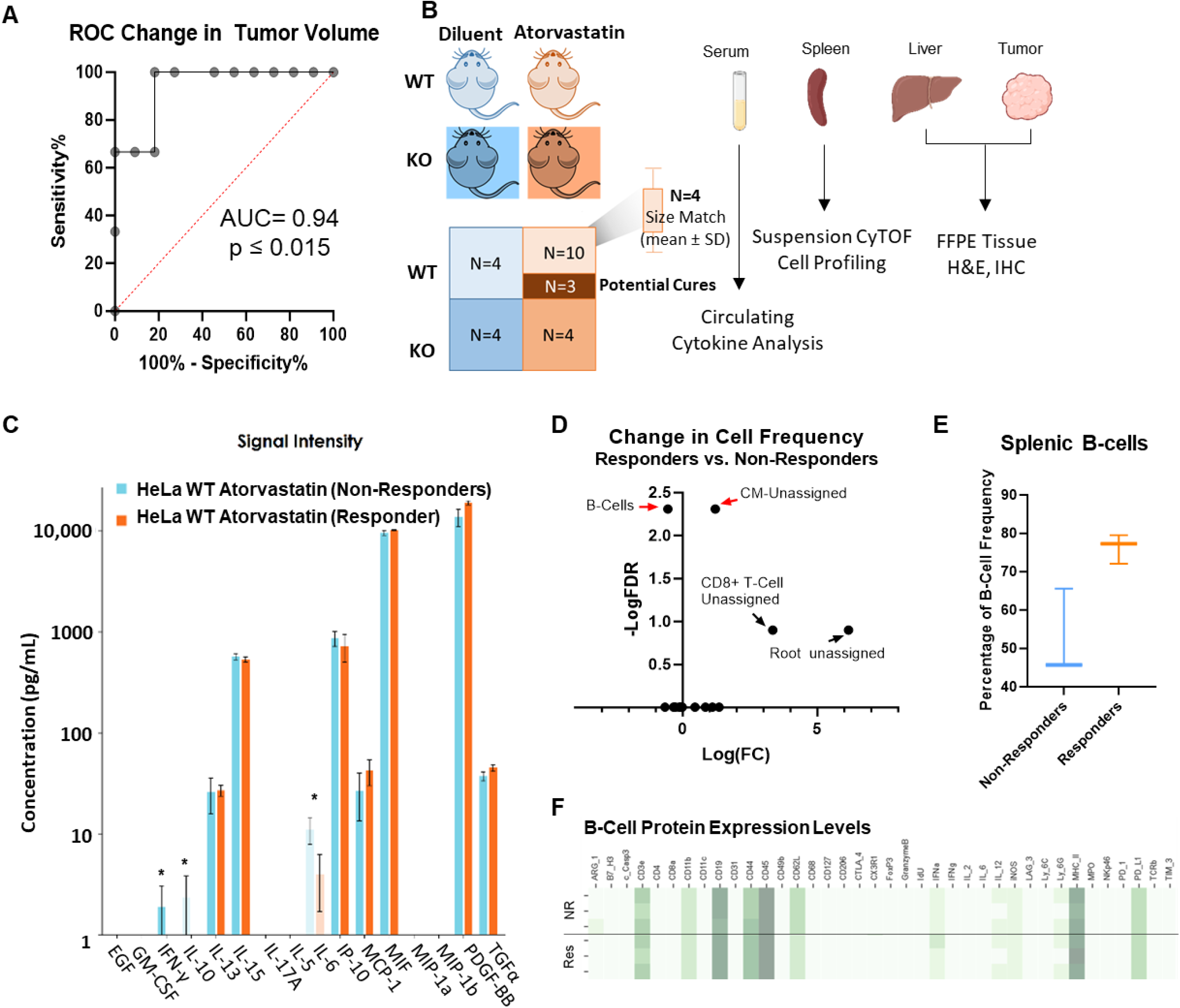
Serum IFN-γ, IL-6 and IL-10 serve as non-invasive PD biomarkers for prevention of tumor outgrowth with atorvastatin treatment. **A.** Receiver operator characteristic curve was generated from previously performed *in vivo* experiments, whereby the change in tumor volume for WT HeLa xenografts treated with atorvastatin was used to predict the fate of the mouse. ROC curves compare the sensitivity and specificity for overall survival by change in tumor volume from week 6 to week 8. Using a cutoff of <0.03 mm^3^, 100% sensitivity was achieved to label the shrinking tumors as “responders” and those that continued to grow as “non-responders.” **B.** Schematic demonstrating the ability to identify potential responders and non-responders to statin treatment for mice bearing WT HeLa xenografts and sacrifice 3-4 mice per treatment condition for downstream analyses to identify biomarkers of response. **C.** Circulating cytokines were measured from plasma collected at experimental endpoint between potential responders and non-responders to atorvastatin treatment. Cytokine concentrations (pg/mL) are represented as mean ± SD for 3 individual mice/group. **D.** Volcano plot highlighting the changes in cell frequency observed following suspension CyTOF analysis of mouse spleens at the designated pre-endpoint between potential responders and non-responders. Statistically significant changes in B-Cell frequency and unassigned cell types were observed. **E.** Box and whisker plot of B-cell frequency between predicted responders and non-responder mice. The box represents the median and first and third quartiles while the whiskers represent the min and maximum for the dataset. 3 mice per group were analyzed. **F.** Heatmap containing protein expression levels for markers used to identify B-cell populations and contained within our panel. Each mouse is represented per row, likely non-responders are grouped on top with potential responders grouped below.

### Patient data supports statin use as a cancer prevention strategy for B7-H3 expressing tumors

Finally, we began to explore the clinical significance of statin-mediated anti-cancer effects in a B7-H3 dependent manner. While 4Ig-B7-H3 is the dominant isoform expressed in human cancers [33], these historical datasets do not explicitly distinguish 2Ig and 4Ig isoforms, and thus, we refer to only B7-H3 in this analysis. Using the publicly accessible TCGA datasets, we performed a tumor wise correlation analysis for CD276 expression across 17 tumor types. For each tumor type, significantly co-expressed genes (p<0.01) were identified and combined into a single matrix containing their Spearman’s correlation coefficient. 22 genes were found to be commonly co-expressed with B7-H3 across all tumor types and this matrix was used to perform pathway enrichment analysis using Enrichr Wiki2019 Pathways [34, 35]. The statin pathway was significantly associated with B7-H3 co-correlated gene sets (p=0.030), as well as others, such as ECM remodeling, MMP function and effects on cell metabolism, and PI3K-Akt signaling **(Figure 7A, Supplemental Table 1)**. To evaluate the potential for B7-H3 to serve as a biomarker of statin efficacy in patient clinical samples, we performed a retrospective chart analysis matched with immunohistochemically stained ovarian cancer tissue specimens. B7-H3 expression was present in both the tumor and stromal compartment (as previously observed) at a high frequency **(Figure 7B, Supplemental Figure 12)**. While expression of B7-H3 in either compartment did not sufficiently separate the overall survival curves to result in a statistically different hazard ratio, patients with higher scores of B7-H3 expression within the tumor (red) trended toward a worse prognosis **(Supplemental Figure 13)**. 269 patient specimens were analyzed, and upon retrospective chart review, 10 patients were prescribed a ‘statin’ at some point in their medical history. Quantitative image analysis of the ovarian tissue specimens was performed and compared between non-statin users (N=259) and statin users (N=10), wherein B7-H3 expression per area of tissue was defined by an H-index and the H-index was compared across the two groups. The mean expression was 88.71 ± 54.13 and 79.33 ± 43.45 for non-statin users and statin users, respectively **(Figure 7C)**. A subset of patients from the study were matched to the cohort of statin-using patients for tumor type, grade and age **(Figure 7D, Supplemental Table 2)** and once matched, their overall survival was compared. Indeed, patients with a history of statin use had significantly (p = 0.05) longer survival than those who were not statin users **(Figure 7E)**. While this sample cohort was too small to compare the dependence of B7-H3 expression within the statin-user cohort, what was striking was the proportion of patients on statins (3.7%), especially given the patient population of post-menopausal women surveyed. This proportion was compared to known general rates of statin use among women in the United States, which is reported to vary from 22% to 28% during the time at which the patient samples were acquired for our retrospective analysis (National Center for Health Statistics. Health, 2010, United States, 2011) [36–39]. A two-tailed one sample proportion test was performed using the conservative estimate of 22% incidence of statin use, wherein the null hypothesis was rejected (p<0.01) **(Figure 7F)**. These data led us to assess the expression of B7-H3 in pre-cancerous lesions across a variety of tumor types. Using tissue microarrays with disease progression tissue specimens (normal, pre-neoplastic disease, neoplasia) for pancreatic, prostate, and breast cancer, immunohistochemical staining revealed increased expression of B7-H3 in pre-neoplastic lesions, including benign prostate hyperplasia (BPH), pancreatic intraepithelial neoplasia (PanIn), and ductal carcinoma in situ (DCIS) **(Figure 7G, Supplemental Figure 14)**. Automated quantification of staining was performed (HistoWiz) and the corresponding H-index was provided for each arrayed tissue spot. A test for trend was performed for each disease site and a significant trend was detected for prostate (***p>0.001), pancreas (*p=0.0461) and breast (*p=0.0104) progression arrays, documenting the clear early presence and increase in B7-H3 expression as tumors develop **(Supplemental Figure 14**). Thus, these data support the clinical utility of systemic cholesterol lowering interventions to complement current cancer treatment strategies and may suggest a powerful role in cancer prevention mechanistically defined by inhibiting 4Ig-B7-H3 dimerization.

**Figure 7.**
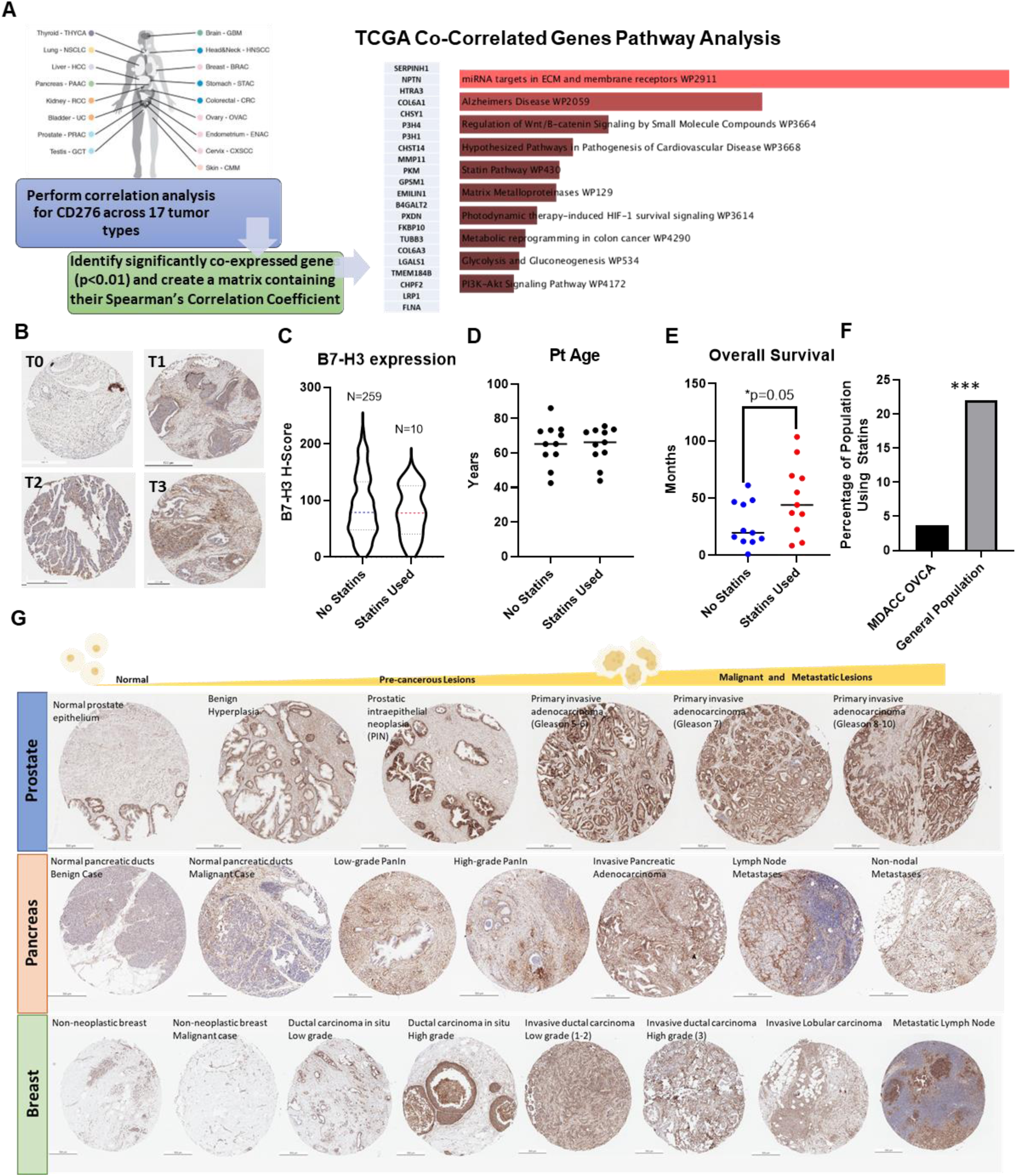
Patient data support statin use as a cancer prevention strategy for B7-H3 expressing tumors. **A.** Schematic demonstrating the workflow used to analyze the publicly available TCGA datasets to identify tumor-wise gene expression correlation analysis for CD276 expression. 17 tumor types were analyzed for the significantly co-expressed genes (p<0.01) and combined into a single matrix containing the Spearman’s Correlation Co-efficient for the commonly co-expressed genes (22 in total). Enrichr pathway analysis was performed and the statin pathway was significantly associated with B7-H3 co-correlated gene sets (p=0.030) as well as others listed in rank order. **B.** Immunohistochemical statin of patient tissue microarrays, representative images for scored tumor 0-3 are depicted. Scale bar represents 250 μm. **C.** 269 patient specimens were analyzed, and upon retrospect chart review 10 patients were taking a ‘statin’ at some point in their medical history. Quantitative image analysis was performed and compared between the non-statin users (N=259) and statin users (N=10), where the B7-H3 expression per area of tissue was defined by an H-index and this was compared across the two groups. The mean expression was 88.71 ± 54.13 and 79.33 ± 43.45 for non-statin users and statin users, respectively. A subset of patients from the study were matched to the cohort of statin using patients for tumor type, grade and age **(D)** and once matched, their overall survival was compared. Patients with a history of statin use had significantly (p = 0.05) longer survival than those who were not statin users **(E)**. **F.** The percentage of patients taking statins among the retrospective chart analysis (3.7%) was plotted and compared with the expected incidence of statin use among post-menopausal women within the United States (22%). A two-tailed one sample proportion test was performed using the estimate of 22% incidence of statin use, where the null hypothesis was rejected (p<0.001). **G.** Immunohistochemical stain of human tissue microarrays showing normal, pre-neoplastic, and neoplastic lesions for several organ types. B7-H3 staining is shown in brown while nuclear stain is shown in blue. Scale bar represents 250 μm.

## Discussion

These studies confirmed the enhanced proliferation and tumorigenic signaling potential of 4Ig-B7-H3 homodimerization/multimerization, wherein inhibition of this protein-protein interaction was associated with decreased tumor cell proliferation, increased immunogenic cell death, and extended survival both *in vitro* and *in vivo*. Statin-mediated disruption of 4Ig-B7-H3 dimerization resulted in both tumor-intrinsic and -extrinsic effects, leading to decreased proliferation and tumorigenic potential. A cholesterol recognition amino acid sequence (CRAC/CARC) was identified within the transmembrane domain of B7-H3 and, as demonstrated through mutagenesis, the contribution of cholesterol-lipid microdomains played a significant role in 4Ig-B7-H3 homodimerization. Decreased clonogenic growth and induction of immunogenic cell death was observed following statin treatment, which translated into functional cures when 4Ig-B7-H3 dimerization was inhibited by pharmacologically relevant doses of statins *in vivo*. Furthermore, pharmacodynamic biomarkers of response were identified when comparing statin responders vs. non-responders *in vivo* and correlated with retrospective clinical data, demonstrating the clinical significance of coupling statins with B7-H3 tumor expression. Thus, our findings support 4Ig-B7-H3 as a clinically relevant biomarker for statin-mediated anti-cancer efficacy.

Our discovery that regulation of 4Ig-B7-H3 dimerization influenced tumor growth has several important implications. These data extend the repertoire of anti-tumor efficacy associated with targeting 4Ig-B7-H3, a promising tumor target given its widespread and differential expression on many solid tumors. Given the surprising yet robust data demonstrating the 4Ig-B7-H3 dependence of statins to inhibit and cure tumor xenografts, these studies also provide evidence that concurrent statin use will alter the baseline dimeric state of 4Ig-B7-H3, which may have implications for other B7-H3-directed therapeutic strategies. While the FLIM data at basal levels of 4Ig-B7-H3 expression reveal 50%/50% monomer/dimer ratios in HeLa cells, hetero-FRET-FLIM experiments with increased levels of 4Ig-B7-H3 in hetero-FRET-FLIM revealed a further shift towards dimerization. These data suggest that expression and other variables (e.g., baseline cholesterol content) may impact the functional state of 4Ig-B7-H3 and thus directly affect 4Ig-B7-H3-directed therapeutic interventions.

In addition to canonical mechanisms of action of statins on cholesterol metabolism, these data provide evidence for a novel mechanism by which statins may afford beneficial effects in the treatment and prevention of cancer, supporting several correlative retrospective clinical studies [40–42]. Our findings suggest a model whereby tumor-specific protein expression of 4Ig-B7-H3 determines the responsiveness to statin treatment. Statins have been shown to alter the immune response, including T-cell signaling, antigen presentation, immune cell migration and cytokine production [43]. The present study raises the possibility that optimal outcomes in cancer prevention and therapy may be achieved utilizing 4Ig-B7-H3 expression as a translatable biomarker to enhance the clinical utility of statin-mediated immune modulation and provide a novel therapeutic approach to reduce the tumorigenic potential of 4Ig-B7-H3. Intriguingly, a significant under representation in statin use among the ovarian cancer patients surveyed at MD Anderson Cancer Center was noted (4%), particularly remarkable given the post-menopausal demographic and widespread statin use (22-28%) among many older adults in the United States. While this may represent a small sample size or could be attributed to lower prescription rates of cholesterol lowering medications for women in the early 2000s (although attempts were made to account for this using historical usage data from the Centers for Disease Control (CDC)), the difference in usage among a population enriched for ovarian cancer patients was striking. Notably, prospective randomized controlled studies are needed before specific recommendations for prevention and treatment can be made; however, the known safety profile and added benefits of statin use among older populations where high cholesterol poses additional health risks for cardiovascular disease, provide support for future studies to prospectively assess the impact of statin use for cancer prevention in high-risk populations and cancer patients with 4Ig-B7-H3-expressing tumors.

In addition, the ability of 4Ig-B7-H3 and other B7-family members to bind cholesterol might be part of the spatiotemporal regulation of these macromolecular complexes necessary for engaging cell-to-cell communication via the immune synapse, which may relate to the immunomodulatory effects observed broadly by cholesterol lowering medications. Therefore, the immune modulatory effects of many anti-inflammatories may broadly be attributed to disruption of protein-protein interactions necessary for engagement between the immune cell and the antigen-presenting cell, as facilitated by cholesterol-mediated super molecular complexes. Although statins provide a safe and effective way to modulate cholesterol-mediated 4Ig-B7-H3 dimerization, our study also demonstrated that the 4Ig-B7-H3 dimerization-inhibitory effect of statin-treatment can be recapitulated by directly disrupting membrane cholesterol levels. Revealing a previously unappreciated target for modulating the tumor immune landscape through disruption of 4Ig-B7-H3 dimerization, and strategies to alter cholesterol metabolism broadly may act synergistically with surgery, chemotherapy or immunotherapy. Similarly, statins have been shown to enhance responses to immune checkpoint therapy (anti-PD-1) in head and neck cancer models [44]. Additionally, other lipid lowering agents, such as aspirin and β-blockers, also enhance responses to PD-1/PD-L1 immune checkpoint therapy for several tumor types, including non-small cell lung cancer, melanoma, and renal cell carcinoma, confirming the enhanced survival observed with baseline statin use [45].

Together, the direct impact of cholesterol-mediated 4Ig-B7-H3 dimerization and the use of statins to influence this cell surface immune modulatory protein-protein interaction may have broad implications in other inflammatory diseases where statin use has a known biological impact and the functional role of 4Ig-B7-H3 expression is underexplored. Of note, this could include cardiovascular disease where patients with high serum levels of B7-H3 had a higher risk of major cardiovascular events [46], yet the functional role of 4Ig-B7-H3 in cardiomyocytes remains unknown. Therefore, there is significant impact and potential for future innovation in exploiting the cholesterol-mediated role of 4Ig-B7-H3 dimerization across fields.

## Methods

### Antibodies and Reagents

Anti-β-actin (4970L), anti-HA-tag (3724S), anti-GFP-tag (2956S), anti-Calreticulin (D3E6) (12238T) antibodies were purchased from Cell Signaling Technology. Anti-GAPDH (G9545) was purchased from Sigma-Aldrich. Anti-B7-H3 antibody was produced in house (MIL33B) or purchased from R&D Systems (AF1397, AF1027). Secondary antibodies were purchased from BioRad Laboratories: goat anti-Mouse (L005680), goat anti-Rabbit (10000045946), rabbit anti-Goat (L006330A). The U2OS-4Ig-B7-H3 cell line was generated using the ReBiL 2.0 plasmid system and cloned using Gibson assembly to express 4Ig-B7-H3 fused with a C-terminal HA-tag linker to N-Luciferase or C-Luciferase. Vector-mEGFP and CD276-mEGFP plasmids were generated by Genecopoeia (Rockville, MD) using CD276 ORF expression clone (NM_00102736.1) or empty control vector for pReceiver-M98 (EX-Neg-M98). Tet-On inducible-homo-dimerization system was purchased from Clontech Laboratories Inc. (Mountain View, California; Catalog: 635068) and 4Ig-B7-H3 was cloned into the construct using PCR-based amplification and In-Fusion assembly according to the manufacturers protocol. CRISPR/Cas9-directed *CD276* knockout HeLa and SKOv3-ip-FLuc cell lines were engineered using a dual plasmid gRNA and Cas9 plasmid system purchased from Santa Cruz Biotechnology and clonally expanded from single cells. Gene knockout was confirmed by qRT-PCR and Western blot analysis.

| Antibody Target | Source | Catalog # | Dilution | Application |
| --- | --- | --- | --- | --- |
| B7-H3 (AF1397) | R&D Systems | AF1397 | 1:2000 | WB |
| B7-H3 (MIL33B) | In house | N/A | 1:1000-5000 | WB |
| GAPDH (G9545) | Sigma Aldrich | G9545-100UL | 1:10,000 | WB |
| B-Actin (13E5) | Cell Signaling Technology | 4970 | 1:10,000 | WB |
| HA-tag (C29F4) | Cell Signaling Technology | 3724 |  |  |
| GFP-tag (D5.1) | Cell Signaling Technology | 2956 |  |  |
| Goat-anti-mouse | BioRad | L005680 | 1:10,000 | WB |
| Goat anti-Rabbit | BioRad | 10000045946 | 1:10,000 | WB |
| Rabbit anti-Goat | BioRad | L006330A | 1:10,000 | WB |
| B7-H3 (AF1397) | R&D Systems | AF1397 | 1:400 | IF, IHC |
| B7-H3 (MIL33B) | In house | N/A | 1:200-500 | IF, IHC |
| B7-H3 (D9M2L) | Cell Signaling Technology | 14058 | 1:100 | IHC |

D-luciferin was purchased from BioGold and prepared as a 30 mg/mL stock solution in phosphate-buffered saline (PBS). Doxycyline was purchased from RPI Research Products International Corp. (Lot 32606) and prepared as a 1mg/mL stock solution in PBS. Methyl-β-cyclodextrin was purchased form Sigma (332615), and Filipin was obtained as part of a cholesterol labeling kit from Abcam (ab133166).Lead compounds were purchased from Selleckchem and prepared as a 10 mM stock solution in DMSO.

Recombinant Human Apolipoprotein A-1 (ApoA1) was purchased from Novus Biologicals (NBP2-34869).

### Cell culture and transfection conditions

Cells were maintained using standard aseptic techniques in a humidified atmosphere of 5% CO_2_. Cell lines were routinely checked for mycoplasma contamination and routine short tandem repeat DNA fingerprinting was performed to ensure proper cell line identification. U2OS ReBiL cell lines were routinely grown in DMEM supplemented with 2mM L-glutamine, nonessential amino acids, and 10% tetracycline-free fetal bovine serum. Gene expression was induced by adding 0.1 µg/mL doxycycline (Dox) as indicated. HeLa and MDA-MB-231 cell lines were grown in DMEM supplemented with 10% FBS and 1% L-Glutamine, SKoV3-ip cell line was grown in McCoy’s 5A, supplemented with 10% FBS and 1% L-Glutamine.

Transient transfection was performed using MegaTran 1.0 (Origene, #TT200005) according to the manufacturer’s protocol. For transfection of a 6-well plate, 3μg of DNA was combined with 9μL of transfection reagent in 250 μL of OPTIMEM and allowed to incubate at room temperature for 10 minutes before being added to the cultured cells. When co-transfection was performed, DNA concentrations were reduced to 1.5 μg of each plasmid.

### Cell proliferation assay

Changes in cancer cell growth with or without B7-H3 expression were determined using clonogenic and sulforhodamine B (SRB) assays. Clonogenic assays were performed using U2OS-4Ig-B7-H3, HeLa (wt, ko, and rescue lines), SKOv3-IP and MDA-MB-231 cell lines, where 2000-5000 cells/well were cultured with or without Doxycycline (if applicable) for 72 hours. Cells were then allowed to grow in a clonogenic fashion for two weeks, and stained with Coomassie blue, before the colonies were quantified. SRB short term cell viability assays were performed using U2OS-4Ig-B7-H3 cells (5×10^4^) cultured with or without doxycycline for 72 hrs. Cells were washed, and fixed with 30% TCA at 4°C, and incubated for 30 min at room temperature with 0.4% SRB in 1% acetic acid. The plates were read with a microplate-reader (Biotek synergy 2, Tecan) at 510 nm.

### Western blotting

Cell lysates were prepared as indicated following incubation in lysis buffer (50 mM Hepes, pH 7.0, 150 mM NaCl, 1.5 mM MgCl_2_, 1 mM EGTA, 10 mM NaF, 10 mM sodium pyrophosphate, 10% glycerol, 1% Triton X-100) plus protease and phosphatase inhibitors (1 mM PMSF, 10 µg/mL leupeptin, 10 µg/mL aprotinin and 1 mM Na_3_VO_4_). Cells were lysed for 30 min on ice, and then centrifuged at 17,000 x g for 30 min at 4°C. The protein concentration was assessed using a bicinchoninic acid (BCA) protein assay (ThermoScientific, #23225). Equal amounts of protein were separated by 8-16% SDS-PAGE, transferred to PVDF membranes and subjected to western blotting using an ECL chemiluminescence reagent (BioRad, #NEL105001).

### ReBiL assay of B7-H3 multimerization

Cells were seeded in 96-well or 384-well black wall, clear bottom plates and treated with doxycycline (100 ng/mL) to induce gene expression. 24-48 hours following gene expression, cells were washed briefly with 1X PBS and D-luciferin (200 µM) was added in phenol red-free DMEM/F12 medium supplemented with MEBSS. Following incubation at 37 °C, bioluminescence was initiated by addition of 100 μL substrate solution containing D-luciferin dissolved in PBS (final concentration = 300 µM). Bioluminescence imaging was performed using an IVIS100 or IVIS Spectrum instrument (FOV C, 1-5 sec exposure, open filter, medium binning). Bioluminescence microscopy of U2OS-4Ig-B7-H3 live cells was performed on a Nikon TiE inverted microscope (Nikon Instruments, Melville, NY, USA) equipped with a back-illuminated 1024 × 1024 pixel CCD with a 13 µm pixel pitch (iKon-M 934; DU934P-BEX2-DD, Andor Inc/Oxford Instruments (Belfast, Northern Ireland), which was air cooled to −85 °C during normal operations with deep depletion fringe suppression and anti-reflective coating [47]. When acquiring bioluminescence images, the camera was read out at 50 kHz, 4X gain, and binning 2 to minimize read noise and maximize sensitivity. Cells were maintained at 37°C in a humid environment for the duration of imaging (10X objective, 20 min acquisition, open filter).

### ReBiL assay of B7-H3 dimerization for high throughput screen

Cells were trypsinized (0.25%), and seeded in white, clear-bottom CELLSTAR μClear® 384-well tissue culture plates (Greiner Bio-One) at 30 000 (3 × 10^4^) cells per well in 40 μL of serum-free, phenol red–free DMEM/F12 medium using the Multidrop Combi (Thermo Fisher). Cells were pre-incubated with doxycycline (final concentration: 500 µg/mL) and D-luciferin (final concentration: 600µM). 48 hours post doxycycline induction of B7-H3 expression, luminescence intensity was measured using a Tecan Infinite M1000. Nanoliter volumes of library compounds or controls (DMSO) are transferred from low dead volume source plates using the LabCyte Echo 550 acoustic transfer platform. The final concentration of DMSO was maintained at 0.3% for all wells excluding those wells containing cells with medium alone. Following 4 h incubation at 37 C, the plates were read a second time and luminescence intensity is determined (Tecan Infinite M1000).

As a counter screen, cell viability analysis using CellTiter-Blue (CTB, Promega) was performed. Immediately following luminescence reading, 10 µL of 1X CTB reagent was dispensed into 384-well plates using the Multidrop Combi; plates were incubated overnight (16 h) at 37 C, and fluorescence was detected using the Tecan Infinite M1000 reader (excitation λ = 560 nm, emission λ = 590 nm). Cell viability was expressed as percent of the mean fluorescent signal intensity of on-plate negative controls. Hit validation was confirmed using a secondary screen with the ReBiL2.0 cell line expressing K-RAS dimerization or n-Luc and c-Luc only cells, and overlapping HITs were removed as potential non-specific targets of the split-luciferase ReBiL 2.0 system, or cell-membrane associated protein complexes.

### Animal models and *in vivo* studies

All animal procedures were approved by The University of Texas M.D. Anderson Cancer Center Animal Study Committee (IACUC). Six week-old athymic nude mice (strain *nu*/*nu*, Charles River) were anesthetized (1.5% isoflurane inhalation) during tumor cell implantation. 5×10^6^ HeLa cervical cancer cells were injected subcutaneously in the rear flank in 100 μL serum-free media. 1×10^6^ SkOv3-ip ovarian cancer cells were injected intraperitoneally in an orthotopic model in 100 μL serum-free media. 1×10^6^ MDA-MB-231 triple negative breast cancer cells were injected into the mammary fat pad beneath the fourth nipple in an orthotopic model in 30 μL 2:1 ratio serum-free media and matrigel. Following xenograft implantation, osmotic pumps (Alzet 1004) were surgically placed subcutaneously containing diluent or atorvastatin (10 mg/kg/day) starting the day of tumor implantation. For HeLa and MDA-MB-231 tumor models, the tumors were measured (l x w) twice weekly using a caliper and the mouse weight was monitored throughout the study. Tumor volume was calculated using V = W^2^×L/2. For the SKOv3-ip model, tumor mass (g) was determined 4 weeks post implantation by excising and pooling each intraperitoneal tumor nodule.

### Immunofluorescent staining

Tumor cells (3×10^4^) were seeded in chamber slides and treated as specified. Cells were fixed with 4% paraformaldehyde and permeabilized with 0.5% Triton X-100. Cells were washed with PBS, and blocked with 5% BSA/PBS followed by incubation with the primary antibody. After washing, cells were incubated with secondary antibodies conjugated with Alexa Fluor 488 or 594 (Molecular Probes, #A11017, #A11020, #A11070, #A11072), mounted and examined using a fluorescence microscope (Nikon TiE microscope) or confocal microscope.

### Fluorescence lifetime imaging analysis

Fluorescence lifetime imaging was performed at the Center for Advanced Microscopy, a Nikon Center of Excellence, Department of Integrative Biology & Pharmacology at McGovern Medical School, UTHealth Houston. Forster resonance energy transfer-fluorescence lifetime imaging microscopy (FRET-FLIM) experiments were performed using the Nikon A1 basic confocal system installed with a PicoQuant FLIM LSM upgrade kit. Data acquisition and analysis were performed using the PicoQuantSymPho Time 32 software. Transiently transfected cells were fixed using 4%PFA at room temperature for 10 minutes and washed briefly with PBS (3X) for 5 minutes and kept shielded from light until analyzed at the Center for Advanced Microscopy. For fluorophore excitation, a 483 nm pulsed laser with an excitation frequency of 40 MHz was applied. This was followed by FLIM measurement of the average amount of time for the fluorophore to remain in the excited state using the PicoQuant SPAD detection unit and FF01 520/35-25 filter. The instrument response function (IRF) was determined at the beginning and end of each imaging session using a Convallaria sample according to the manufacturers protocol. Photon count rate was kept under 10% of the pulse rate, and enough frames were acquired to obtain the cumulative signal intensity of at least 10^4^ photons. Fluorescence decay curves were fitted to a bi-exponential reconvolution function adjusted to the IRF and the average lifetime was calculated and represented in the FLIM images as τ. The lifetime of 4Ig-B7-H3-mEGFP was measured and compared to Vector-mEGFP, following adjustment of transfection conditions such that each protein was expressed at similar levels as determined by both Western blot analysis and surveying the imaging field with epifluorescence microscopy. All measurements were taken from whole-field images of cells, and at least 10 measurements were taken for each analysis over 3 biological replicates.

### Circulating Cytokine expression

For animals enrolled in the responder vs. non-responder study, serum was obtained via cardiac puncture, used with the IsoPlexis single-cell highly multiplexed proteomic platform for cytokine analysis. 2 uL of serum from each animal was used in technical duplicate with the CODEPLEX Innate Immune Secretome panel (Panel-2L 12-2) for murine species, following the manufacturers protocol. Cytokine levels were analyzed using the IsoSpeak2.9.0 software.

### Suspension CyTOF

Table of antibodies used for CyTOF shown in **Supplemental Table 3**. HeLa tumor-bearing WT and *CD276^−/−^* animals with and without statin treatment were processed for CyTOF according to the pre-determined ROC-derived metrics of responders and non-responders (when tumor volume decreased over two measurements, at day 50-56). Animals were then euthanized by carbon dioxide asphyxiation. Single-cell suspension of cells were harvested from tumors, spleen, and liver. Samples were processed for CyTOF according to conventional protocols including barcoding and staining extracellular proteins followed by permeabilization and intracellular proteins. CyTOF data were analyzed using Phenograph and Astrolabe [48]. For Phenograph analysis, clusters with no marker expression above median were excluded. Myeloid cell populations identified by Phenograph and Astrolabe are consistent with those previously reported [49].

### Immunohistochemistry and survival analysis

Immunohistochemical staining was performed by Histowiz (B7-H3) or RHCL (Immune cell infiltrates) according to established protocols. Survival analysis was performed following retrospective chart review under IRB PA17-0959.

### Statistical Analysis

Data are expressed as mean ± standard deviation, unless otherwise specified. Statistical significance was analyzed using a two-tailed students *t* test, Wilcoxon-rank log test, or One-way and two-way ANOVA for multiple group comparisons. P-values <0.05 were considered statistically significant and denoted by a single asterisk. P-values <0.01 were denoted by a double asterisk. Experiments were performed in biological triplicate (three independent experiments conducted over independent timeframes) unless otherwise specified and often included multiple technical replicates in each experiment (same conditions within a given experiment). Specific experimental details are included in the figure legends.

Statistical parameters of assay performance were calculated according to the following formulas:

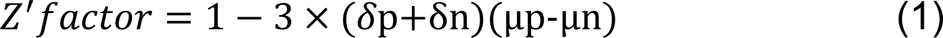

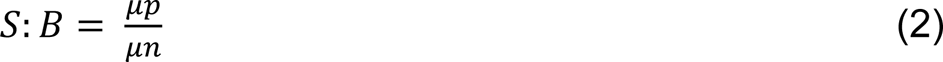

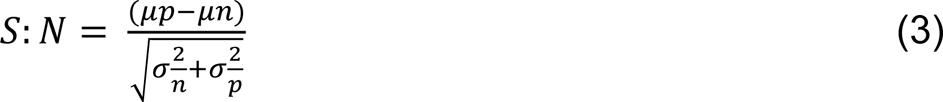

where δ_p_ and δ_n_ are standard deviation of the positive control group p and the negative control group n, and μ_p_ and μ_n_ are the arithmetic means of the two groups, respectively; S:B, signal to background; S:N, and signal-to-noise [50]. For cell-based assays, a Z′ of ≥0.5 signifies that outliers can be reliably identified as statistically significant despite well-to-well and plate-to-plate variability. We calculated Z′ using Eq. 1, which is based on the mean and standard deviation of positive and negative controls and is calculated in the absence of library compounds [51, 52]. Based on this equation, Z′ is improved by greater signal separation between the mean of positive and negative controls, as well as by reducing variance between replicates (i.e., standard deviation). In practical terms, consistency between replicates would improve confidence in a single well outlier being truly significant (i.e., the compound treatment in that single well resulted in significant changes in complex formation, rather than the change in luminescence being due to simple well-to-well variability).

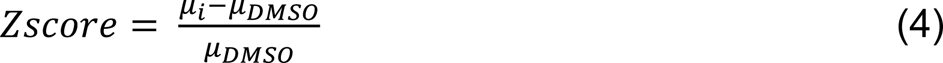

where μ_i_ and μ_DMSO_ are the arithmetic means of the sample (i.e., screened compound) and 0.3% DMSO control group, respectively. Robust Z and Z’ were calculated as described above substituting the arithmetic median instead of the mean for each variable.

### Data Availability

Source data is contained within the manuscript and supplemental information.

## Acknowledgements

The authors would like to thank members of the D. Piwnica-Worms laboratory for discussion and suggestions, Dr. Geoffrey Wahl and Yao-Chang Li for providing the ReBiL2.0 platform constructs and parental U2OS cell line, and Dr. Reid Powell and Ivy Nguyen at the Texas A&M HTSMSC for their help with running the high throughput screen. Additionally, this work would not have been possible without the expertise and assistance of many shared resource core facilities, including the Research Histology Core Lab (MDACC), the Flow Cytometry Core Facility (MDACC), the Small Molecule Screening Core (Texas A&M IBT), and the UTHealth Center for Advanced Microscopy, a Nikon Center of Excellence, Department of Integrative Biology & Pharmacology at McGovern Medical School, UTHealth Houston. The RHCL and FCCF at MDACC obtain support from NCI P30 CA016672. The SMSC at Texas A&M IBT receives support from the Cancer Prevention & Research Institute of Texas grant number RP200668.

This work was supported by NIH Grant P50 CA94056 to the MD Anderson Cancer Center– Washington University Inter-Institutional Molecular Imaging Center; the Gerald Dewey Dodd, Jr., Endowed Distinguished Chair of the University of Texas MD Anderson Cancer Center; and a generous gift from the Estate of Barbara Cox Anthony/Koch Foundation. M.N.S. was supported by a Ruth L. Kirchstein Postdoctoral Individual Research Service Award (F32) NCI F32CA250323, the Harold C. and Mary L. Dailey Endowed Fellowship Recognition of Research Excellence (M.D. Anderson Cancer Center), the Diane Denson Tobola Endowed Fellowship in Ovarian Cancer Research (M.D. Anderson Cancer Center), the Quantitative Imaging Analysis Core via the QIAC Partnership in Research Pilot Project Program at The University of Texas MD Anderson Cancer Center, and support from the Brain SPORE (2P50CA127001) and Ovarian Cancer (P50CA281701) SPORE CEP programs.

