## Supplemental Information for "Statins inhibit onco-dimerization of the 4Ig isoform of B7-H3"

**Author Contributions:** M.N.S. S.T.G. and D.P.W. designed the study. M.N.S., S.E.G., A.A, A.N. and P.Y. carried out experiments. D.P.W. provided resources and reagents. J.L. provided TMA and pathology consult, P.B. provided clinical consult, M.N.S., S.T.G., S.E.G., A.A., and D.P.W. analyzed data, and M.N.S. and D.P.W. wrote the manuscript. All authors edited and approved the manuscript.

**Competing Interest Statement:** The authors declare no competing interests. The University of Texas MD Anderson Cancer Center has filed a patent application on an anti-B7-H3 antibody used in this report (S.T.G., and D.P.W., inventors). The technology is licensed in part to Radiopharm Ventures, LLC. The remaining authors have no conflicts of interest.

**Keywords:** B7-H3, CD276, protein dimerization, immune checkpoint, split-luciferase complementation, statins, immune modulation, gynecological cancers

*Corresponding author:
David Piwnica-Worms, M.D., Ph.D.
1881 East Road, Unit 1907
Houston, TX  77054
Email address:


**Supplemental Figure 1**

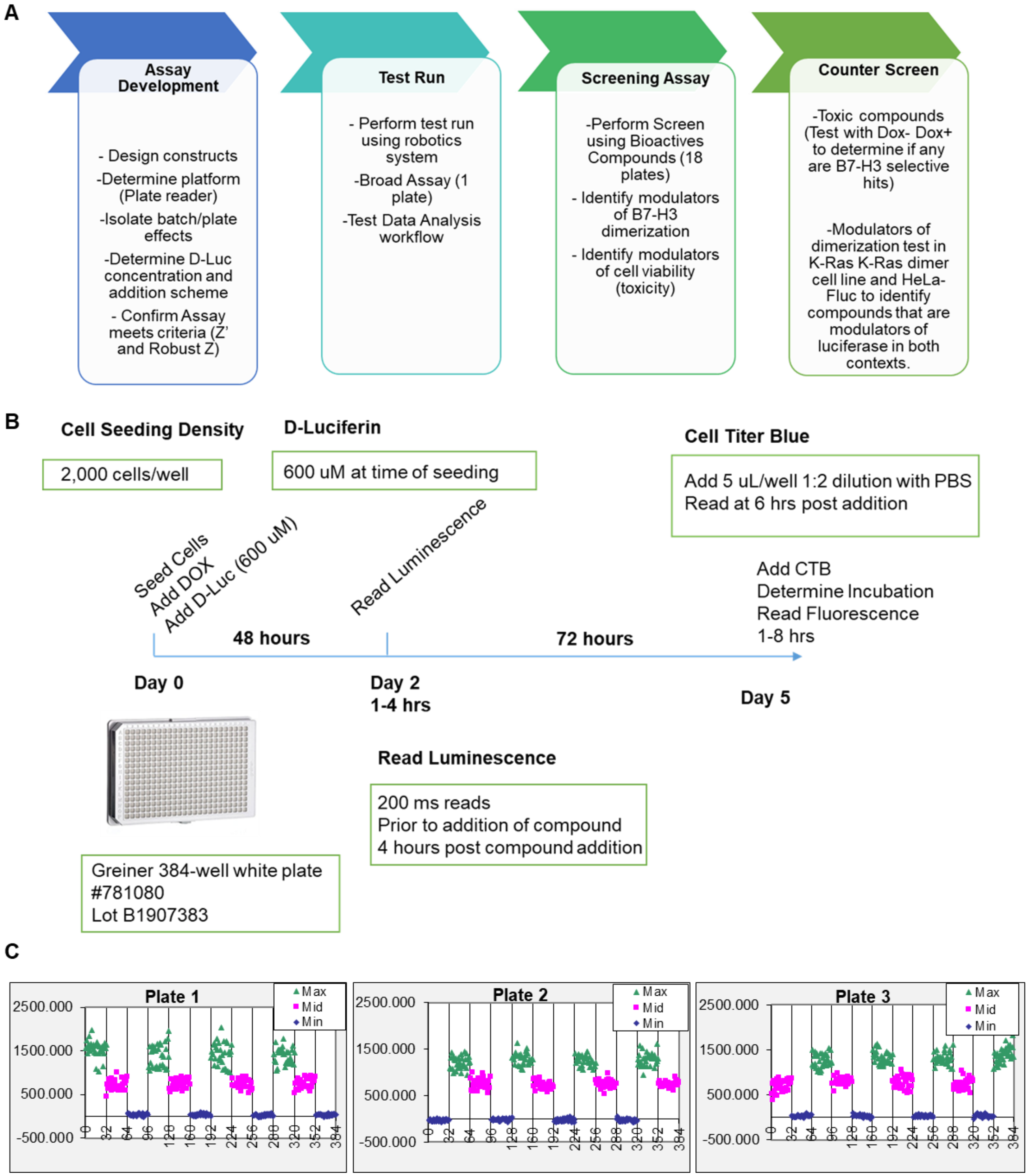

**Supplemental Figure 1. High throughput screening conditions. A.** Schematic of the screening process is outlined taking into account the criteria being tested at each step to develop the HTS. **B.** Assay specific conditions and methods are detailed for each plate. **C.** Luminescence readouts for interleaved-signal format plates using a min, mid and max signal concentration of doxycycline to induce 4Ig-B7-H3 expression and dimerization as observed by split-luciferase complementation.

**Supplemental Figure 2**

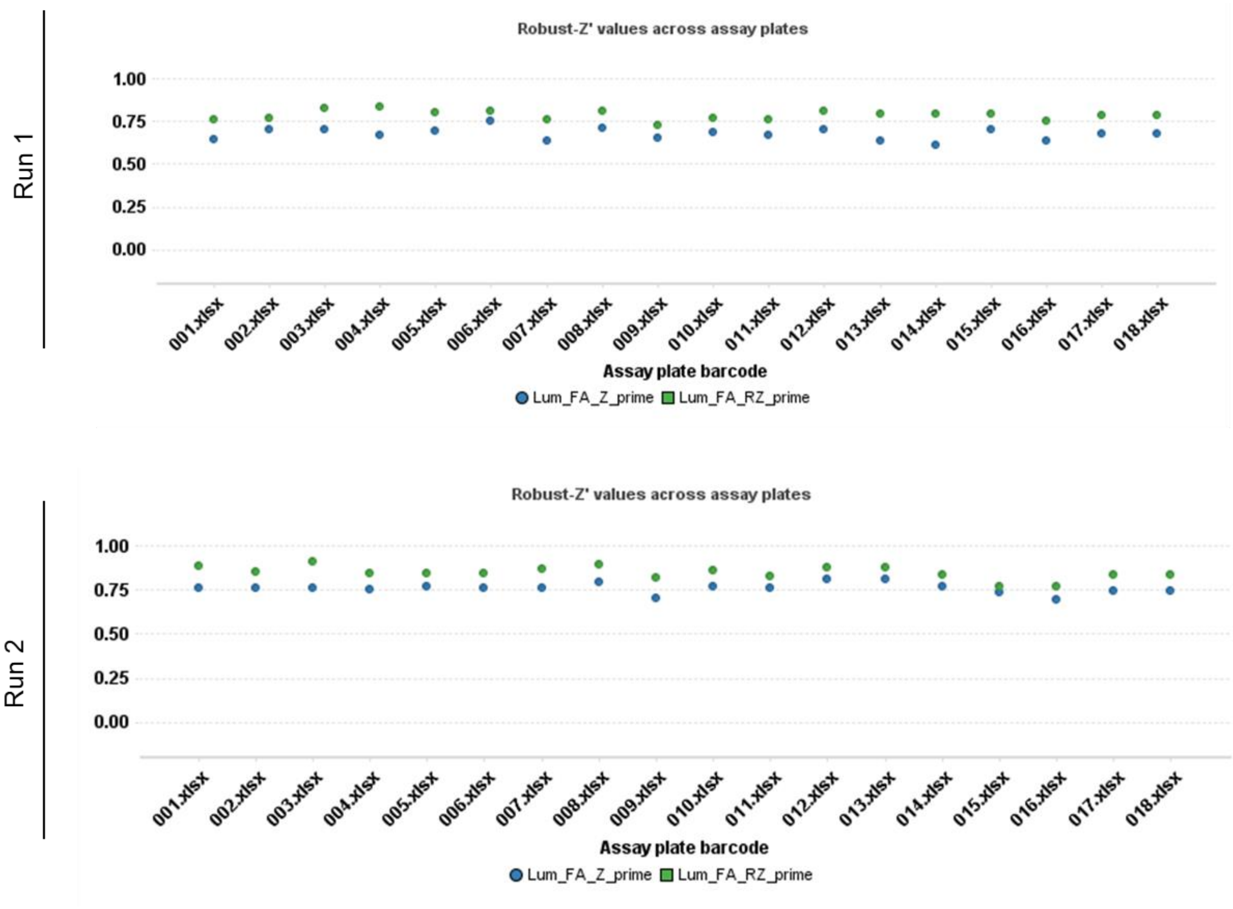

**Supplemental Figure 2. The high-throughput screen was robust across all plates and biological replicates.** Z' and Robust Z' values for each plate were calculated and plotted across biological replicates (Run 1 and Run 2) of the HTS.

**Supplemental Figure 3**

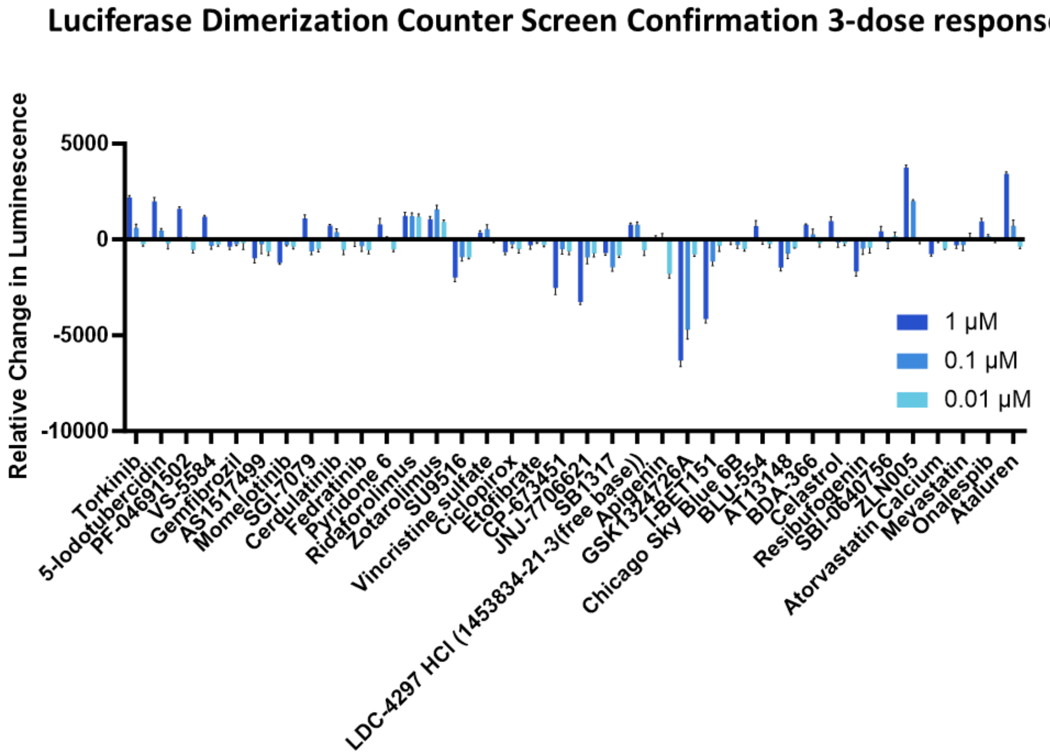

**Supplemental Figure 3. 4Ig-B7-H3 dimerization counter screen was performed using a 3-dose titration curve for all "active compounds".** U2OS-4Ig-B7-H3 cells were treated with active compounds for 4 hrs at 0.1 -1.0 μM concentrations. Relative change in luminescence is plotted where bars represent the mean ± SD. The experiment was performed in technical triplicate.

**Supplemental Figure 4**

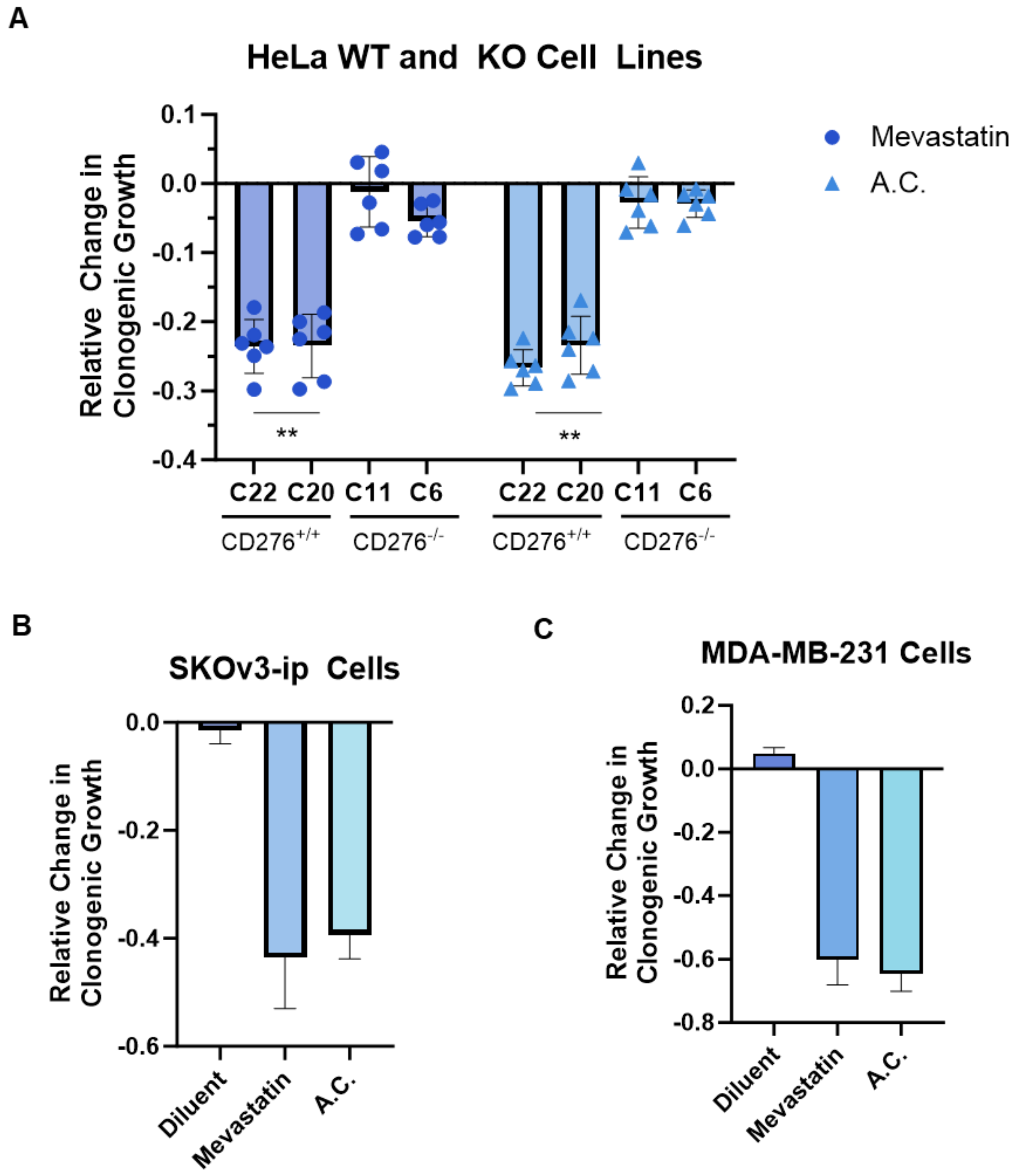

**Supplemental Figure 4. Statins inhibitors reduce clonogenic growth in multiple 4Ig-B7-H3 positive cancer cell lines. A.** 2,000 HeLa WT or KO cells were seeded and without statin treatment (1µM treatment with mevastatin, atorvastatin calcium) on day 1 and day 4 and clonogenic growth was measured after 10 days. Colonies were fixed and stained with Coomassie blue on day 10. Clonogenic growth was quantified using ImageJ and represented in the bar graph (mean ± SD). The experiment was performed in technical triplicate with three biological replicates, ***p<0.001. **B-C.** Clonogenic growth was also measured for SKOv3-ip (5,000 cells/well seeded) and MDA-MB-231 (2,000 cells/well).

**Supplemental Figure 5**

```
B7-H4  ------------------------------------------------------------    0
B7-H3  RSPTGAVEVQVPEDPVVALVGTDATLRCSFSPEPGFSLAQLNLIWQLTDTKQ--LVHSFT    298
PD-L1  WHLLNAFTVTVPKDLYVVEYGSNMTIECKFPVEKQLDLAALIVYWEMEDKNIIQFVHGEE    72
PD-L2  HQIAALFTVTVPKELYIIEHGSNVTLECNFDTGSHVNLGAITASLQ--------------    60

B7-H4  ------------------MASLGQILFWSIISIIIILAGAIALIIGFGI-----------    31
B7-H3  EGRDQGSAYANRTALFPDLLAQGNASL-RLQRVRVADEGSFTCFVSIRDF-GSAAVSLQV    356
PD-L1  DLKVQHSSYRQRARLLKDQLSLGNAAL-QITDVKLQDAGVYRCMISYGGA-DYKRITVKV    130
PD-L2  KVENDTSPHRERATLLEEQLPLGKASF-HIPQVQVRDEGQYQCIIIYGVAWDYKYLTLKV    119
                             *:  :  :  : :   *    ::

B7-H4  ----SAFSMPEVNVD--YNASSETLRCEAPRWFPQPTVVWASQVDQGANFSEVSNTSFEL    85
B7-H3  AAPYSKPSMTLEPNKDLRPGDTVTITCSSYRGYPEAEVFWQDGQGV--PLTGNVTTSQMA    414
PD-L1  NAPYNKINQRILVVD--PVTSEHELTCQ-AEGYPKAEVIWTSSDHQ--VLSGKTTTTNSK    185
PD-L2  KASYRKINTHILKV---PETDEVELTCQ-ATGYPLAEVSWPNVSVP-------ANTSHSR    168
              .            .   : *.    :*   * * .            .*:

B7-H4  NSENVTMKVVSV-LYNVTINNTYSCMIENDIAKATGDIKVTESEIKRR------SHLQLL    138
B7-H3  NEQGL-FDVHSVLRVVLGANGTYSCLVRNPVLQQDAHGSVTITGQPMTFPPEALWVTVGL    473
PD-L1  REEKL-FNVTSTLRINTTTNEIFYCTFRRLDPEENHTAELVIPELPLAHPPNERTHLVIL    244
PD-L2  TPEGL-YQVTSVLRLKPPPGRNFSCVFWNTHVREL-----TLASIDLQSQMEPRTHPTWL    222
         : :  .* *.       .  : * . .   .       .                  *

B7-H4  NSKASLCVSSFF-AISWAL-LPLSPYLMLK------------------------------    166
B7-H3  S-VCLIALLVALAFVCWRK----IKQSCEEEN-AGAEDQDGEGEGSKTALQPLKHSDSKE    527
PD-L1  G-AILLCLGVALTFIFR-----LRKGRMMDVKKCGIQDTNSK-----------KQSDTHL    287
PD-L2  L-HIFI-PSCIIAFIFIATVIALRKQLCQ--KLYSSKDTTKR-----------PVTTTKR    267
            :     :  :

B7-H4  ------- 166
B7-H3  DDGQEIA 534
PD-L1  EET---- 290
PD-L2  EVNSAI- 273
```

**Supplemental Figure 5. CRAC/CARC domains are present in the transmembrane region of many B7-family members.** Sequence alignment of B7-H4, B7-H3, PD-L1 and PD-L2 highlight the presence of predicted CRAC/CARC domains within all four B7-family member proteins. The stars, dots and colons below the alignment indicate degree of conservation. “*” indicate columns of identical residues, colons and dots indicate columns where there is some conservation of the biochemical character of the side chains.

**Supplemental Figure 6**

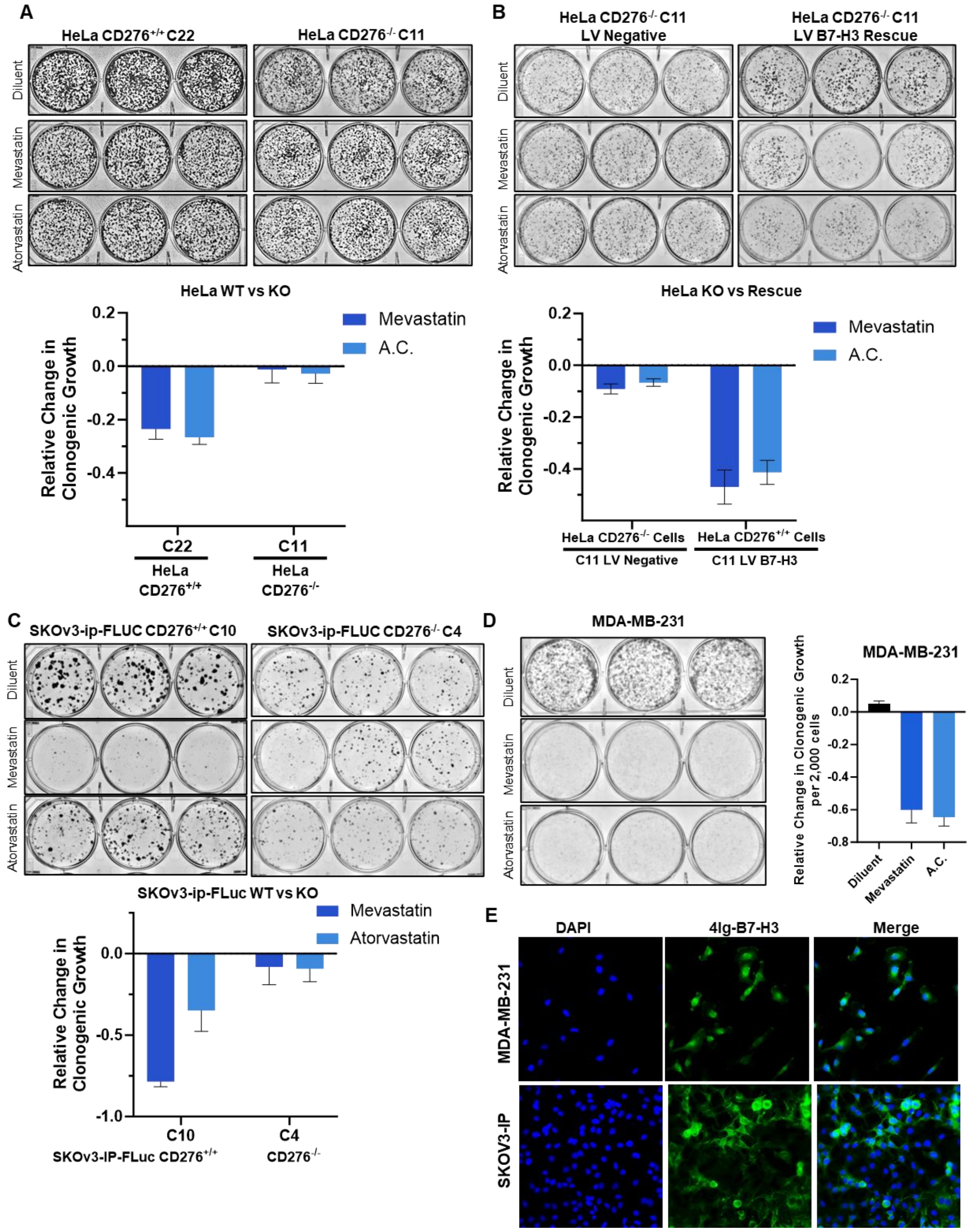

**Supplemental Figure 6. Statins inhibit clonogenic growth in a 4Ig-B7-H3 dependent fashion. A-B.** 2,000 HeLa wildtype or *CD276* knockout cells, or rescued cells **(B)** were seeded and allowed to grow for 10 days, changing the media every 72 hours. Cells were subjected to 1µM treatment with mevastatin or atorvastatin calcium on day 1 and day 4. Plates were stained with Coomassie blue dye on day 10 and representative images of the plates are shown. **C.** 5,000 SKOv3-ip wildtype or *CD276* knockout cells were seeded and allowed to grow for 10 days, changing the media every 72 hours. Cells were subjected to 1µM treatment with mevastatin or atorvastatin calcium on day 1 and day 4. Plates were stained with Coomassie blue dye on day 10 and representative images of the plates are shown. **A-C.** Clonogenic growth was quantified using ImageJ and represented in the bar graph (mean ± SD). The experiment was performed in technical triplicate with three biological replicates, ***$p<0.001$, significance was reached when comparing each individual treatment between the B7-H3 expressing group and the non-expressing group (KO). **D.** 2,000 MDA-MB-231 WT cells were seeded and allowed to grow for 10 days, changing the media every 72 hours. Cells were subjected to 1µM treatment with mevastatin or atorvastatin calcium on day 1 and day 4. Plates were stained with Coomassie blue dye on day 10 and representative images of the plates are shown. Clonogenic growth was quantified using ImageJ and represented in the bar graph (mean ± SD). The experiment was performed in technical triplicate with three biological replicates, ***$p<0.001$. **E.** Endogenous expression of B7-H3 as determined by immunofluorescence staining using SKOv3-ip and MDA-MB-231 cancer cells.

### Supplemental Figure 7

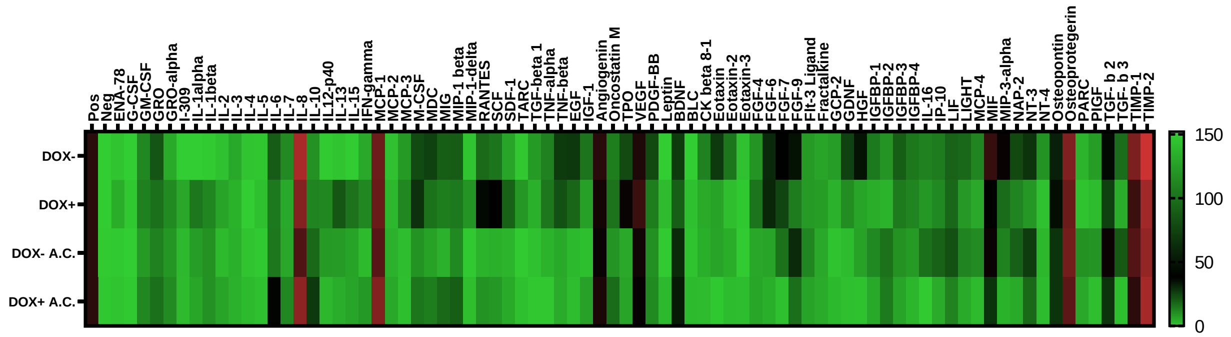

**Supplemental Figure 7. Statin treatment alters cytokine release, some of which occur in a 4Ig-B7-H3-dependent fashion.** Heatmap cytokine protein levels are plotted for media collected from 1.0 x $10^5$ U2OS-4Ig-B7-H3 cells seeded and treated with or without Doxycycline for 48 hours followed by treatment with diluent or atorvastatin (1 μM) for 24 hours. Relative cytokine levels were plotted for each condition; DOX-, DOX+, DOX- Atorvastatin treatment, DOX+ Atorvastatin treatment.

**Supplemental Figure 8**

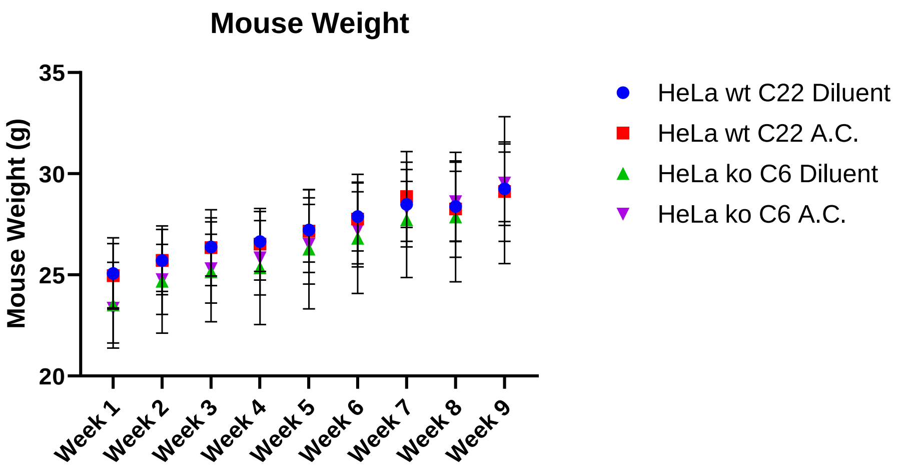

**Supplemental Figure 8. Treatment of HeLa WT or HeLa KO xenografts with statins or diluent did not significantly alter mouse weight over the course of the study.** Mouse weight for each mouse was measured weekly, mean ± SD, for each treatment group is presented.

### Supplemental Figure 9

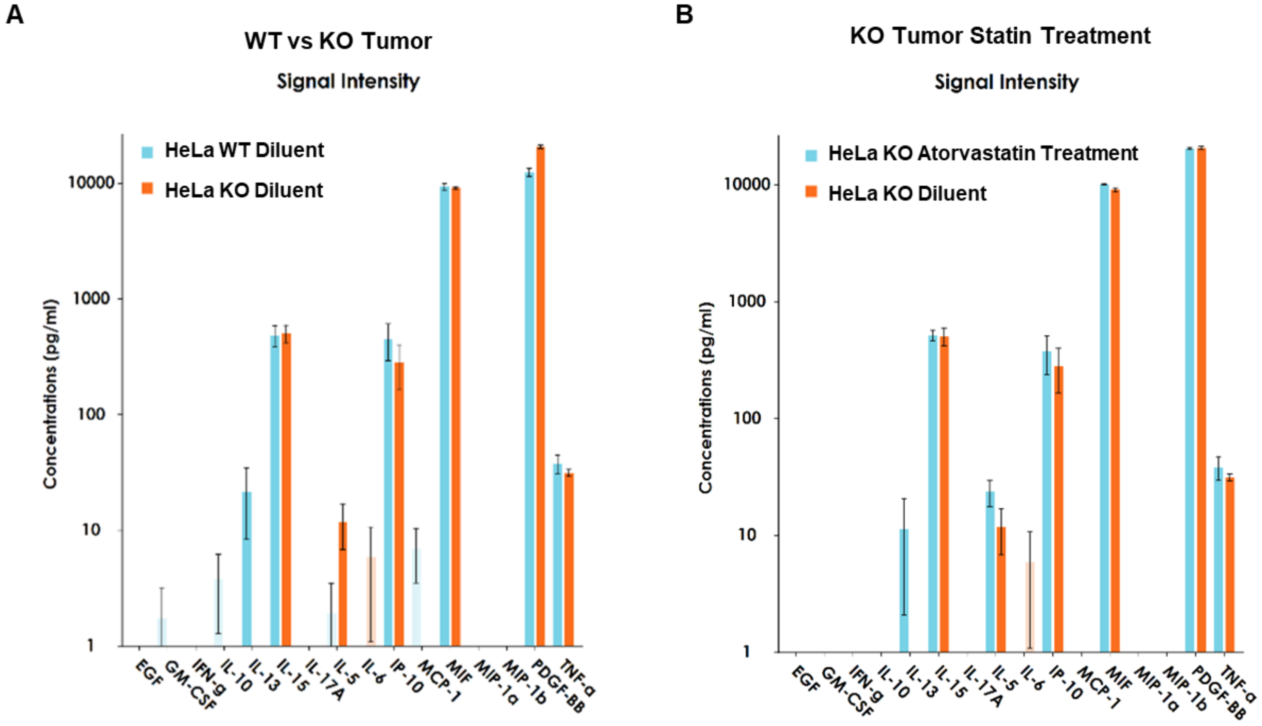

**Supplemental Figure 9. Serum cytokine levels for *in vivo* tumor types (WT and KO) and response to statin treatment.** Circulating cytokines were measured from plasma collected at experimental pre-endpoint, where mice bearing WT or KO HeLa xenografts were treated with diluent or atorvastatin (as described in Figure 6). Cytokine concentrations (pg/mL) are represented as mean ± SD for 3 individual mice/group.

**Supplemental Figure 10**

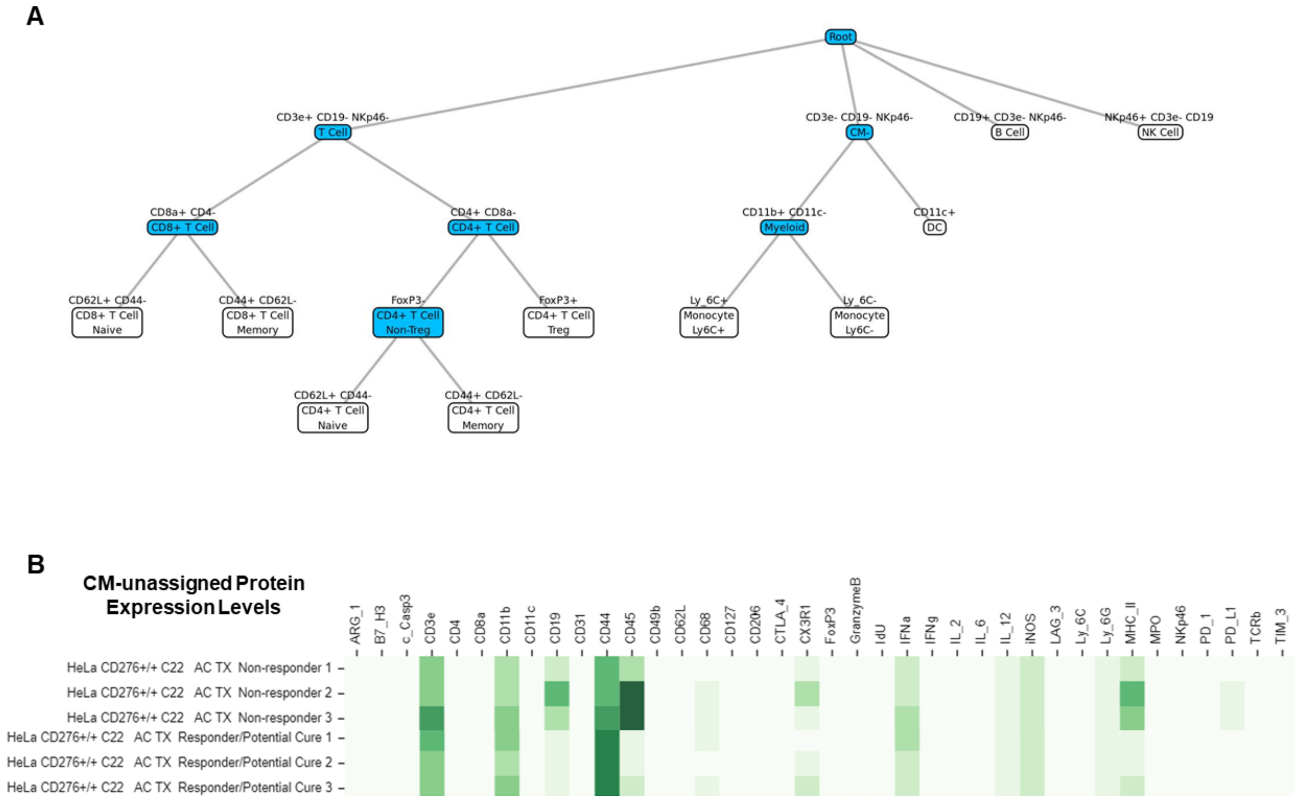

**Supplemental Figure 10. Astrolabe hierarchy of cell identification and protein content within the CM-unassigned cell class. A.** Schematic documenting Astrolabe hierarchy of cell identification markers. **B.** Heatmap containing protein content levels for markers within the CM-Unassigned cell populations contained within our panel. Each mouse is represented per row; likely non-responders are grouped on top with potential responders grouped below.

**Supplemental Figure 11.**

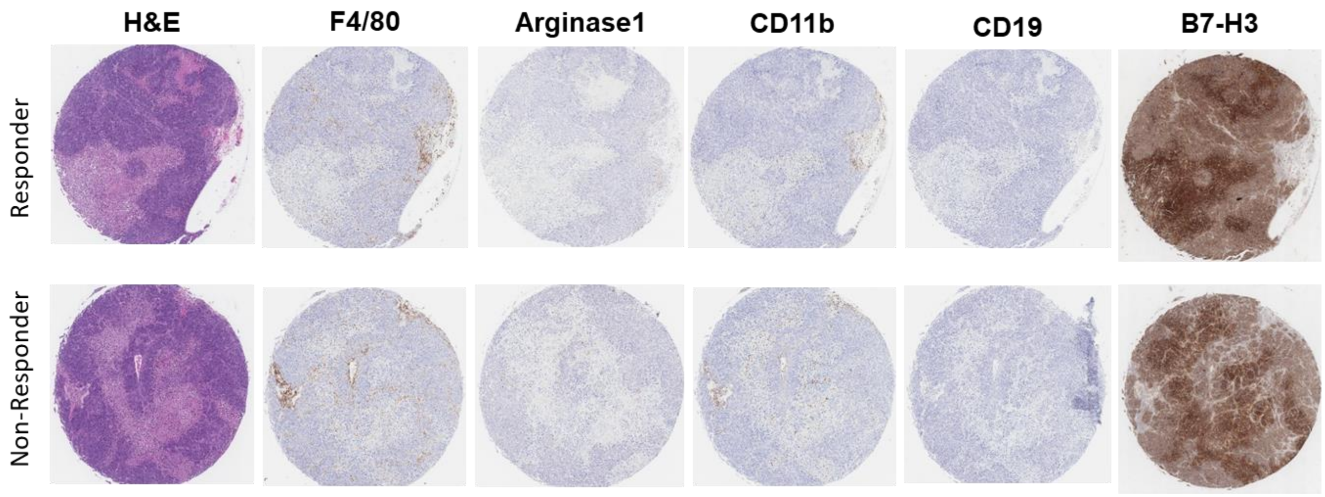

**Supplemental Figure 11. Immunohistochemical staining of tumor immune infiltrates in the tumor of responders vs. non-responders to atorvastatin treatment.**

**Supplemental Figure 12**

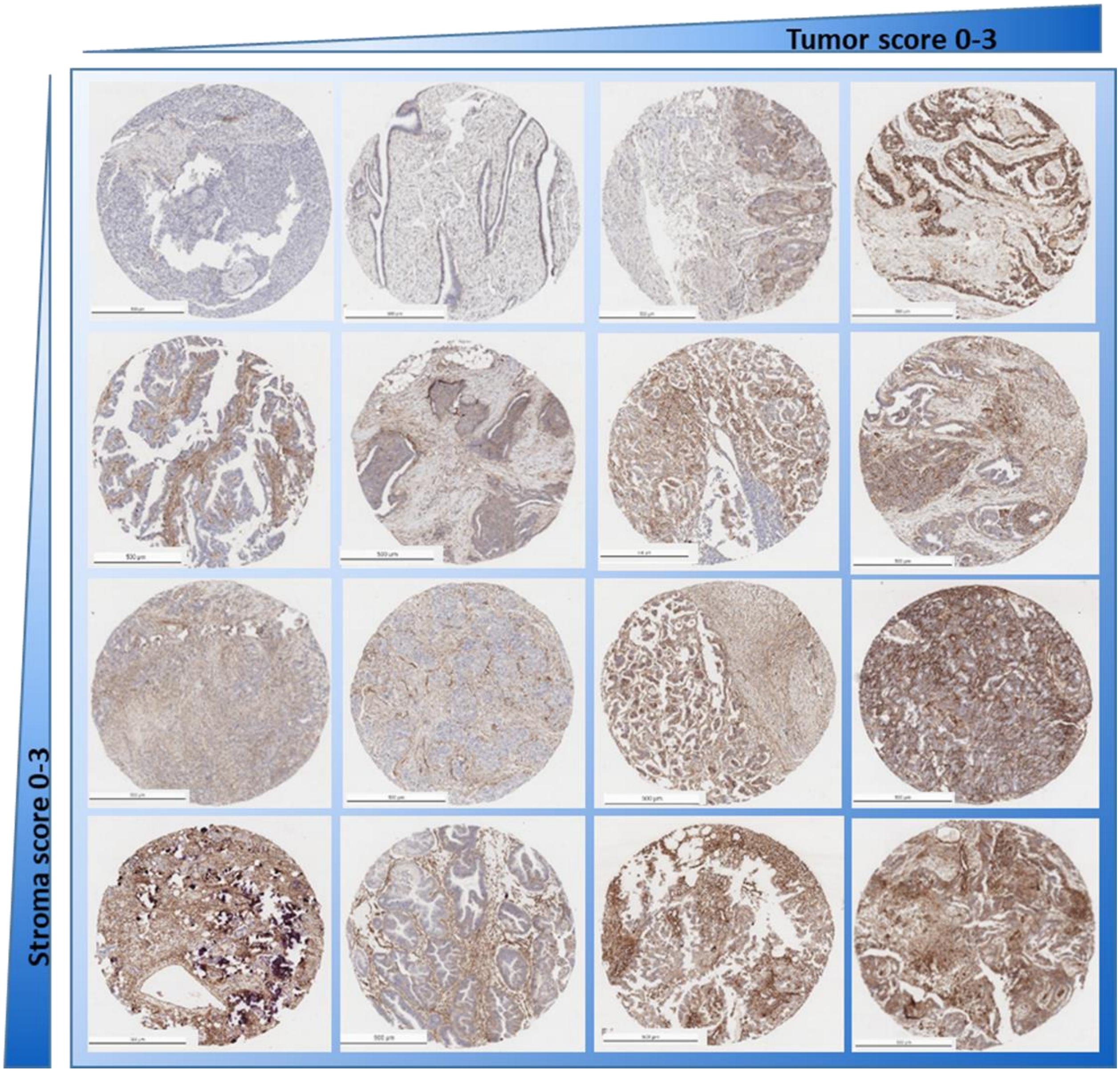

**Supplemental Figure 12.** Immunohistochemical staining for B7-H3 of patient tissue microarrays, representative images for scored tumor and stroma from 0-3 are depicted. Scale bar represents 250 µm.

**Supplemental Figure 13**

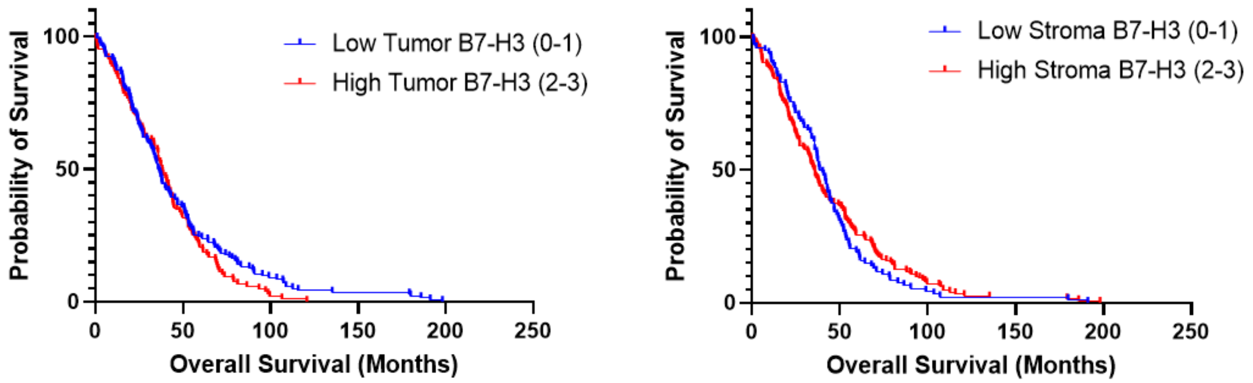

**Supplemental Figure 13. Kaplan-Meier curves for patient survival based on tumor and stromal staining scores of B7-H3 protein.**

### Supplemental Figure 14

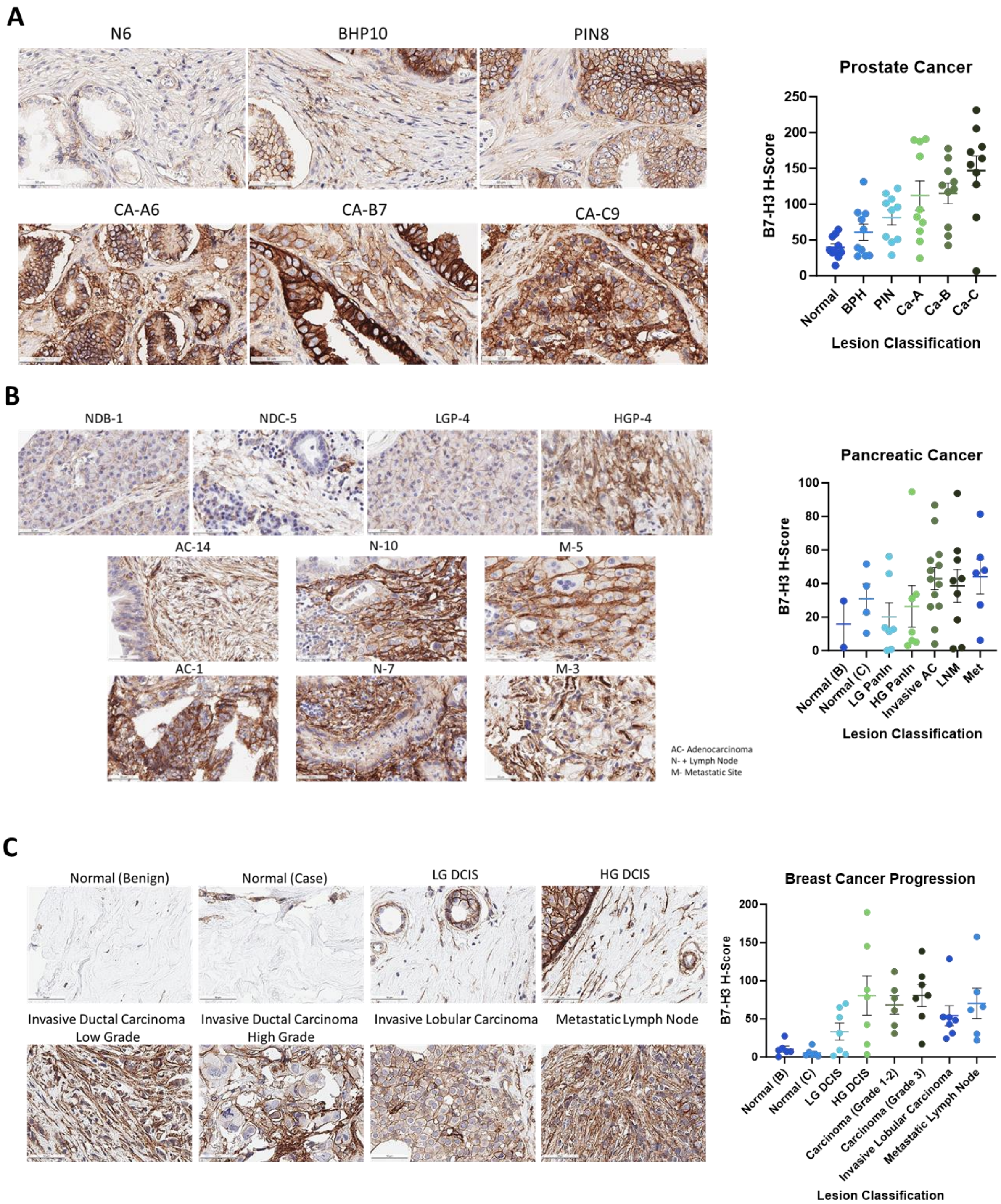

**Supplemental Figure 14.** Immunohistochemical staining of B7-H3 of patient tissue microarrays for tumor progression across multiple disease sites. Representative images of B7-H3 staining across progression of disease and corresponding H-Score per arrayed tissue spot as determined by automated quantification of the staining for prostate cancer (A), pancreatic cancer (B), and breast cancer (C). Scale bar represents 50 μm.

**Supplemental Table 1.** Wiki 2019 Pathway Analysis list of p-Values for by term for pathway enrichment.

| Term | P-value |
|---|---|
| miRNA targets in ECM and membrane receptors WP2911 | 0.000239501 |
| Alzheimer's disease WP2059 | 0.003394807 |
| Regulation of Wnt/B-catenin Signaling by Small Molecule Compounds WP3664 | 0.017707727 |
| Hypothesized Pathways in Pathogenesis of Cardiovascular Disease WP3668 | 0.025937094 |
| Statin inhibition of cholesterol production WP430 | 0.030027116 |
| Matrix Metalloproteinases WP129 | 0.031047063 |
| Photodynamic therapy-induced HIF-1 survival signaling WP3614 | 0.038158147 |
| Metabolic reprogramming in colon cancer WP4290 | 0.043207035 |
| Glycolysis and Gluconeogenesis WP534 | 0.046224242 |
| PI3K-Akt signaling pathway WP4172 | 0.048926293 |

**Supplemental Table 2. Clinicopathological Variables for Ovarian Cancer TMA**

| Clincopathological variable | | n |
|---|---|---|
| Histologic types | | |
| Low-grade serous carcinoma | | 22 |
| High-grade serous carcinoma | | 250 |
| Other/Unknown | | 12 |
| Age at diagnosis (years) | | (STD) |
| Mean | 60.45 | 12.3 |
| Median | 61.2 | |
| Range | 21.7-92.4 | |
| FIGO Stage | | |
| Stage I | | 18 |
| Stage II | | 11 |
| Stage III | | 182 |
| Stage IV | | 62 |
| Unknown | | 11 |
| Diagnosis | | |
| Clear Cell | | 8 |
| Endocervical | | 2 |
| Endometrial Papillary Serous Carcinoma | | 1 |
| Endometrioid Adenocarcinoma | | 15 |
| Malignant Mixed Mullerian Tumor (MMMT) | | 6 |
| Mixed Type Carcinoma | | 38 |
| Mucinous | | 12 |
| Serous Carcinoma | | 189 |
| Transitional Cell Carcinoma | | 5 |
| Undifferentiated Carcinoma | | 8 |

**Supplemental Table 3.** CyTOF panel antibodies.

| TaggedAbs.description | Target | Label | IntracellularStaining | Clone | Specificities | Source | Catalog |
|---|---|---|---|---|---|---|---|
| Ly-6G 141Pr | Ly-6G | 141Pr | FALSE | 1A8 | Ms | DVS-Fluidigm | 3141008B |
| Caspase 3(Cleaved) 142Nd | Caspase 3, cleaved | 142Nd | TRUE | D3E9 | Hu, Rt, Ms, crossed | DVS-Fluidigm | 3142004A |
| TCRb(Ms) 143Nd (MDA) | TCRbeta | 143Nd | FALSE | H57-597 | Ms | Tonbo | 70-5961-U100 |
| IL-2(Ms) 144Nd | IL-2 | 144Nd | TRUE | JES6-5H4 | Ms | DVS-Fluidigm | 3144002B |
| CD274(Ms) 145Nd | CD274, PD-L1, B7-H1 | 145Nd | FALSE | 10F.9G2 | Ms | Tonbo | 70-1243-U100 |
| CD8a(Ms) 146Nd (MDA) | CD8a | 146Nd | FALSE | 53-6.7 | Ms | BioLegend | 100702 |
| CD223(Ms) 147Sm | CD223, LAG-3 | 147Sm | FALSE | C9B7W | Ms | BioLegend | 125202 |
| CD11b 148Nd | CD11b | 148Nd | FALSE | M1/70 | Ms, Hu | DVS-Fluidigm | 3148003B |
| CD19(Ms) 149Sm (MDA) | CD19 | 149Sm | FALSE | 4D5 | Ms | BioLegend | 115502 |
| Ly-6C 150Nd | Ly-6C | 150Nd | FALSE | HK1.4 | Ms | DVS-Fluidigm | 3150010B |
| CD335(Ms) 151Eu | CD335, NKp46 | 151Eu | FALSE | 29A1.4 | Ms | BioLegend | 137625 |
| CD3e 153Eu | CD3e | 153Eu | FALSE | D4V8L | Ms | CST | 99940BF |
| IFNa(Ms) 154Sm | IFNa | 154Sm | TRUE | RMMA-1 | Ms | PBL | 22100-1 |
| MPO 155Gd | MPO | 155Gd | TRUE | EPR20257 | Hu, Ms, Rt | Abcam | ab221847 |
| CD4(Ms) 156Gd | CD4 | 156Gd | FALSE | GK1.5 | Ms | Tonbo | 70-0041-U100 |
| Foxp3(Ms) 158Gd | Foxp3 | 158Gd | TRUE | FJK-16s | Ms, Rt, Bv, Cn, Po, Fe | DVS-Fluidigm | 3158003A |
| IL-12(Ms) 159Tb | IL-12 | 159Tb | TRUE | C15.6 | Ms | BD | 554477 |
| CD44 160Gd | CD44 | 160Gd | FALSE | IM7 | Hu, Ms, Ch, Rh | BioLegend | 103002 |
| iNOS(Ms) 161Dy | iNOS | 161Dy | TRUE | CXNFT | Ms | DVS-Fluidigm | 3161011B |
| TIM-3(Ms) 162Dy | TIM3, CD366 | 162Dy | FALSE | RMT3-23 | Ms | DVS-Fluidigm | 3162029B |
| CD152(Ms) 163Dy | CD152, CTLA-4 | 163Dy | FALSE | 9H10 | Ms | BioLegend | 106202 |
| CD62L(Ms) 164Dy | CD62L | 164Dy | FALSE | MEL-14 | Ms | DVS-Fluidigm | 3164003B |
| IFNg(Ms) 165Ho | IFNg | 165Ho | TRUE | XMG1.2 | Ms | DVS-Fluidigm | 3165003B |
| Arginase-1 166Er | Arginase-1 | 166Er | TRUE | Polyclonal | Hu, Ms | DVS-Fluidigm | 3166023B |
| IL-6(Ms) 167Er | IL-6 | 167Er | TRUE | MP5-20F3 | Ms | DVS-Fluidigm | 3167003B |
| CD206(Ms) 169Tm | CD206, MMR | 169Tm | TRUE | C068C2 | Ms | DVS-Fluidigm | 3169021B |
| CD49b(Ms) 170Er | CD49b, Integrin Î±2 | 170Er | FALSE | HMÎ±2 | Ms | DVS-Fluidigm | 3170008B |
| CD279(Ms) 171Yb (29F.1A12) | CD279, PD-1 | 171Yb | FALSE | 29F.1A12 | Ms | BioLegend | 135202 |
| Granzyme B 173Yb | Granzyme B | 173Yb | TRUE | GB11 | Hu, Ms | DVS-Fluidigm | 3173006B |
| I-A/I-E(Ms) 174Yb | I-A/I-E, MHC-II | 174Yb | FALSE | M5/114.15.2 | Ms | DVS-Fluidigm | 3209006B |
| CD127(Ms) 175Lu | CD127, IL-7Ra (Ms) | 175Lu | FALSE | A7R34 | Ms | DVS-Fluidigm | 3175006B |
| CX3CR1(Ms) 176Yb | CX3CR1 | 176Yb | FALSE | SA011F11<br>SA011F11<br>SA011F11 | Ms | BioLegend | 149002 |
| CD11c(Ms) 209Bi | CD11c | 209Bi | FALSE | N418 | Ms | DVS-Fluidigm | 3209005B |
| CD45(Ms) 89Y | CD45(Ms) | 89Y | FALSE | 30-F11 | Ms | DVS-Fluidigm | 3089005B |
| B7-H3 | CD276 (B7-H3) | | FALSE | AF1397 | Hu, Ms | R&D Systems | |

**Supplemental Table 4.** HTS Screening Parameters

| Category | Parameter | Description |
|---|---|---|
| Assay | Type of assay | Split Luciferase Complementation, Cell Viability |
| | Target | 4Ig-B7-H3 homodimerization |
| | Primary measurement | Change in luminescence |
| | Key reagents | D-Luc, Doxycycline, |
| | Assay protocol | 30,000 cells were pretreated with doxycycline (500 µg/mL) and D-luciferin (600µM) for 48 hours. Luminescence was read using a Tecan Infinite M1000, prior to the addition of compounds in nanoliter volumes using the LabCyte Echo 550 acoustic transfer platform. Following 4 hr incubation at 37°C, luminescence was re-read. Final cell viability was measured 72 hours post compound addition using CellTiter-Blue (Promega) fluorescence assay. |
| | Additional comments | Final DMSO concentrations were maintained at 0.3% for all wells excluding those containing cells with medium alone |
| Library | Library size | 5362 compounds |
| | Library composition | Small molecules with |
| | Source | Texas A&M IBT High Throughput Screening Core |
| | Additional comments | |
| Screen | Format | 384-well plate |
| | Concentration(s) tested | 1.0 µM |
| | Plate controls | DMSO, DOX-, DOX+ |
| | Reagent/ compound dispensing system | LabCyte Echo 550 |
| | Detection instrument and software | Tecan Infinite M1000, Magellan software |
| | Assay validation/QC | Assay QC was performed using interleaved-signal format to assess inter- and intra-assay variation over multiple days. Coefficients of variation were calculated for each signal on each plate and all found to be less than 20% Signal windows and Z' factors for each plate were ≥ 2 or ≥ 0.4, respectively.<br><br>Screen was performed in biological duplicate, a counter screen was performed to eliminate inhibitors of split luciferase complementation. Validation was performed in an orthogonal luminescence readout, and using time- and dose-dependent |
| | Correction factors | Removal of compounds where the variation in pre-read between duplicate assays was ≥ 10% |
| | Normalization | Robust Z-scores were used to normalize the change in luminescence across plates for untreated, and DMSO control wells. |
| | Additional comments | Active compounds were further refined to "On-target" compounds following the counter screen using K-Ras split-luciferase ReBil 2.0 assay. |
| Post-HTS analysis | Hit criteria | Change in Robust Z-score |
| | Hit rate | 0.65% |
| | Additional assay(s) | CellTiter-Blue for cell viability and general toxicity |
| | Confirmation of hit purity and structure | Replicate assays were confirmed using commercially purchased compounds, validated through NRM |
| | Additional comments | Secondary assays (Homo- and HeteroFRET-FLIM) were performed to confirm the inhibition of 4Ig-B7-H3 dimerization with statins. |
